# Prediction of residue-specific contributions to binding and thermal stability using yeast surface display

**DOI:** 10.1101/2021.05.31.446445

**Authors:** Shahbaz Ahmed, Munmun Bhasin, Kavyashree Manjunath, Raghavan Varadarajan

**Author notes:** Author for correspondence: -. 2612, **Email:**.

## Abstract

Accurate prediction of residue burial as well as quantitative prediction of residue-specific contributions to protein stability and activity is challenging, especially in the absence of experimental structural information. This is important for prediction and understanding of disease causing mutations, and for protein stabilization and design. Using yeast surface display of a saturation mutagenesis library of the bacterial toxin CcdB, we probe the relationship between ligand binding and expression level of displayed protein, with *in vivo* solubility in *E.coli* and *in vitro* thermal stability. We find that both the stability and solubility correlate well with the total amount of active protein on the yeast cell surface but not with total amount of expressed protein. We coupled FACS and deep sequencing to reconstruct the binding and expression mean fluorescent intensity of each mutant. The reconstructed mean fluorescence intensity (MFI_seq_) was used to differentiate between buried site, exposed non active-site and exposed active-site positions with high accuracy. The MFI_seq_ was also used as a criterion to identify destabilized as well as stabilized mutants in the library, and to predict the melting temperatures of destabilized mutants. These predictions were experimentally validated and were more accurate than those of various computational predictors. The approach was extended to successfully identify buried and active-site residues in the receptor binding domain of the spike protein of SARS-CoV-2, suggesting it has general applicability.

## Introduction

Mutagenesis is often used to generate variants of proteins with improved biophysical properties such as solubility and activity and to understand protein function. The advancement of high-throughput mutagenesis techniques has enabled the generation of a large number of variants of a protein in a short span of time, in a massively parallelizable manner (Jain & Varadarajan, 2014; Zheng *et al*, 2004; Wrenbeck *et al*, 2016). If an appropriate functional assay to score protein activity *in vivo* exist, it is possible to infer the relative activity of each variant in the library, through library screening coupled to next generation sequencing (Adkar *et al*, 2012; Fowler *et al*, 2010; Matreyek *et al*, 2018). However, there is a dearth of efficient, high-throughput methods to measure the solubility and stability of multiple protein variants in parallel, and to discriminate between buried and active-site residues solely using mutational data (Bhasin & Varadarajan, 2021).

Yeast surface display (YSD) is commonly used as a tool to identify protein variants with improved biophysical properties (Jones *et al*, 2006; Schweickhardt *et al*, 2003). YSD is preferable to bacterial expression for disulfide containing or glycosylated proteins. Agglutinin based Aga2p is the most widely used system to display proteins on the yeast cell surface (Shusta *et al*, 2008). Aga2p is a small protein (7.5 kDa), covalently linked via disulphide linkages to the yeast cell surface protein Aga1p (Boder & Wittrup, 1997). Previous studies have shown that the amount of protein displayed on the yeast cell surface is directly correlated to the amount of protein secreted by the cells, as well as the thermal stability of the protein (Shusta *et al*, 1999). However, in other studies where the secretion efficiency (Hagihara & Kim, 2002) or yeast cell surface expression of proteins was measured, no such correlation was observed (Park *et al*, 2006a; Piatesi *et al*, 2006). Proteolysis of yeast surface displayed proteins has also been used to differentiate properly folded, stable variants from unstructured variants or molten globules, as a proxy for stabilization (Chevalier *et al*, 2017; Rocklin *et al*, 2017; Basanta *et al*, 2020). However, this has primarily been applied to relatively small proteins (Chevalier *et al*, 2017; Rocklin *et al*, 2017; Basanta *et al*, 2020; Dou *et al*, 2018)

A previous study which showed correlation between stability and expression levels was carried out on a limited number of mutants, that were studied individually. In addition, the WT protein itself had a very low T_m_ (Shusta *et al*, 1999). It has also been suggested that if the stability of a protein crosses a certain threshold, its expression does not increase linearly with increase in stability and it is therefore difficult to distinguish stable mutants from less stable ones, using only expression as the criterion (Traxlmayr & Shusta, 2017). With a very high level of yeast surface expression for unstable variants, the yeast quality control system may not be able to differentiate between properly folded, unfolded or molten globule like proteins. However, once displayed on the yeast cell surface such mutants may unfold or aggregate and hence will not bind to a tertiary structure specific ligand or cognate partner.

To verify the above hypothesis, we used *Escherichia.coli* (*E.coli*) CcdB as a model protein. CcdB is the toxin component of the CcdAB toxin-antitoxin (TA) module which binds both free DNA Gyrase and the DNA Gyrase-DNA complex, these are referred to as inhibition and poisoning respectively. Formation of the poisoned CcdB:DNA Gyrase:DNA ternary complex stalls replication and causes cell death (Bernard & Couturier, 1992). The other component of this TA module codes for an antitoxin CcdA, which neutralizes the toxicity of the CcdB toxin upon binding to CcdB. A mutation of Arginine to Cysteine in the DNA Gyrase subunit A (GyrA) at residue 462 can abolish the binding of Gyrase to CcdB (Bernard & Couturier, 1992). The CSH501 *E.coli* strain carries this mutation in the gene of the *gyrA* subunit which makes it insensitive to CcdB (Bajaj *et al*, 2008a). In a previous study, a single-site saturation mutagenesis library of CcdB was generated and the mutants were scored based on their *in vivo* growth phenotype (MS_seq_ score) (Adkar *et al*, 2012). In *E.coli*, a good correlation was found between the MS_seq_ score of ∼70 mutants with either ΔT_m_ of purified protein (r =0.65) or *in vivo* solubility in *E.coli* (r =0.69) (Tripathi *et al*, 2016). In contrast to plate based phenotypes, YSD provides greater flexibility and improved quantitation. We therefore wished to explore the correlation between the amount of surface expression or ligand binding seen with YSD, with thermal stability and *E.coli in vivo* solubility using this large set of characterized mutants, which had a range of *in vitro* thermal stability and *in vivo* solubility.

We initially examined 30 different variants of CcdB, which have varying solubility (when expressed in *E.coli)*, *in vitro* thermal stability, accessibility and residue depth. The *in vivo* solubility of these mutants ranged from completely soluble to insoluble. We did not find a good correlation between total expressed protein amount on the yeast cell surface and either *in vivo* solubility in *E.coli,* or *in vitro* determined thermal stability. However, a better correlation was observed between the amount of active protein on the yeast cell surface (i.e., the amount of bound ligand) with *in vivo* solubility/thermal stability. In the yeast cell surface display system (Chao *et al*, 2006), activity was monitored by measuring the extent of binding of yeast cell surface displayed CcdB to a FLAG tagged fragment of GyrA14 as described previously (Sahoo *et al*, 2015).

Multiple rounds of sorting enrich mutants which have the highest expression and binding on the yeast cell surface. Sorting in such a way may lead to the identification of mutants with better biophysical properties, however, it does not give any information about the relative activity of all the mutants in a library. We coupled FACS and deep sequencing to reconstruct the MFI (MFI_seq_) of each mutant in the Site Saturation Mutagenesis (SSM) library of CcdB, using single round FACS sorting methodology. We use this parameter MFI_seq_, to rank all the mutants based on their activity to generate the mutational landscape or distribution of fitness effects (DFE). We found that the DFE generated using binding was more accurate than the DFE generated using expression. Overall, our MFI_seq_ scoring parameter could readily discriminate between stable and destabilized mutants of CcdB in a highly multiplexed manner.

It is well known that mutations that affect activity occur primarily at either surface exposed residues directly involved in binding or catalysis or at buried residues important for folding and stability. It has been difficult to distinguish between these two classes of residues, solely from mutational data (Bhasin & Varadarajan, 2021). We show here that by examining the effects of charged substitution on surface expression we can discriminate between the two classes of residues. To further validate the approach described above, we analyzed previously published saturation mutagenesis YSD expression and binding data for the receptor binding domain (RBD) of SARS-CoV-2 to its ligand ACE-2 (Starr *et al*, 2020). We could successfully predict both binding-site and buried residues solely from the mutational data in this system as well.

## Results

### YSD of CcdB mutants

Yeast surface display (YSD) has become an increasingly popular tool for protein engineering and library screening applications (Pepper *et al*, 2008). Aga2p mating adhesion receptor of *Saccharomyces cerevisiae* is used as a fusion protein for yeast surface display. For surface expression, we used a vector in which CcdB is fused at the C-terminus of Aga2 (Sahoo *et al*, 2015). We generated (Supplementary Figure S1) and individually characterized 30 CcdB variants on the yeast cell surface. Most CcdB mutants had similar levels of expression to the WT protein (Figure 1A). However, the mutants showed different amounts of active protein as assayed by binding to the FLAG tagged GyrA14 compared to the WT protein (Figure 1B). Previously, we have characterized the *in vitro* thermal stability and *in vivo* solubility of several CcdB mutants (Tripathi *et al*, 2016). The amounts of total and active protein were estimated using antibodies against the HA-tag at the N-terminal of the yeast surface displayed CcdB and the C-terminal FLAG tag of GyrA14 respectively. The correlation coefficient (r) between amount of total protein on the yeast cell surface with *in vivo* solubility or T_m_ of the corresponding purified protein were 0.59 and 0.56 respectively (Figure 2A-B). It is unclear why mutants which have very low solubility in *E.coli* are highly expressed on the yeast cell surface. It was previously hypothesized that the protein folding quality control system in yeast is not as effective as in mammalian systems, therefore partially folded/molten globule/aggregated protein may exist on the surface of yeast (Park *et al*, 2006a). A correlation of r=0.85 was found between the amount of active protein on the yeast cell surface with its *in vivo* solubility determined in *E.coli* (Figure 2C). We also found a better correlation (r=0.80) between amount of active CcdB protein on the yeast cell surface and its *in vitro* thermal stability (Figure 2D), compared to that between total CcdB protein on the yeast cell surface and thermal stability.

**Figure 1:**
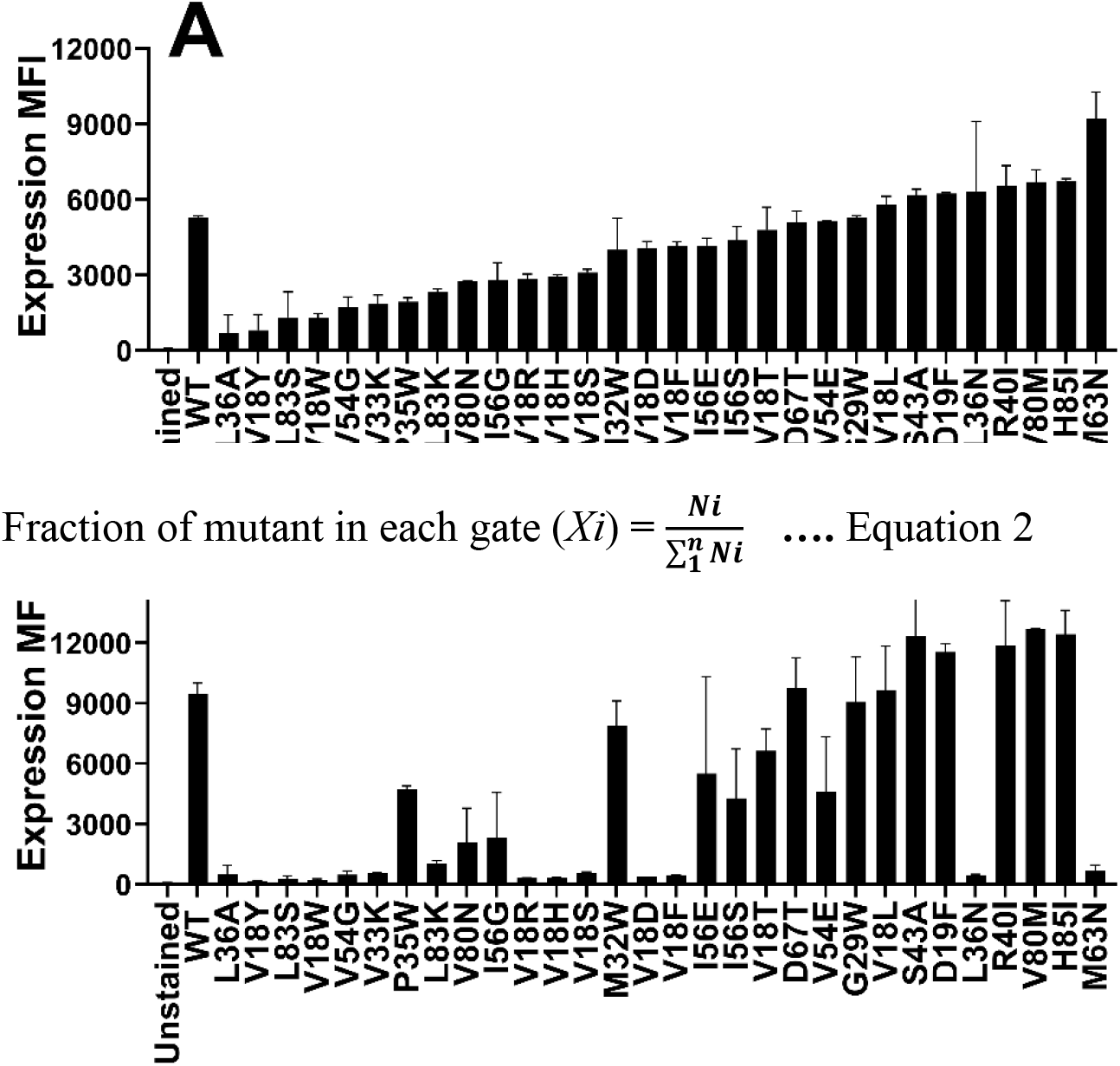
Comparison of the level of expression and binding of CcdB mutants on the yeast cell surface. (A) The expression and (B) binding to GyrA14 of individual mutants. Errors are calculated from two biological replicates. Most mutants expressed at high levels, however, the amount of active protein varied widely. A few mutants which showed a high level of expression did not show any binding to GyrA14. In both panels, mutants are arranged in order of increasing expression level.

**Figure 2:**
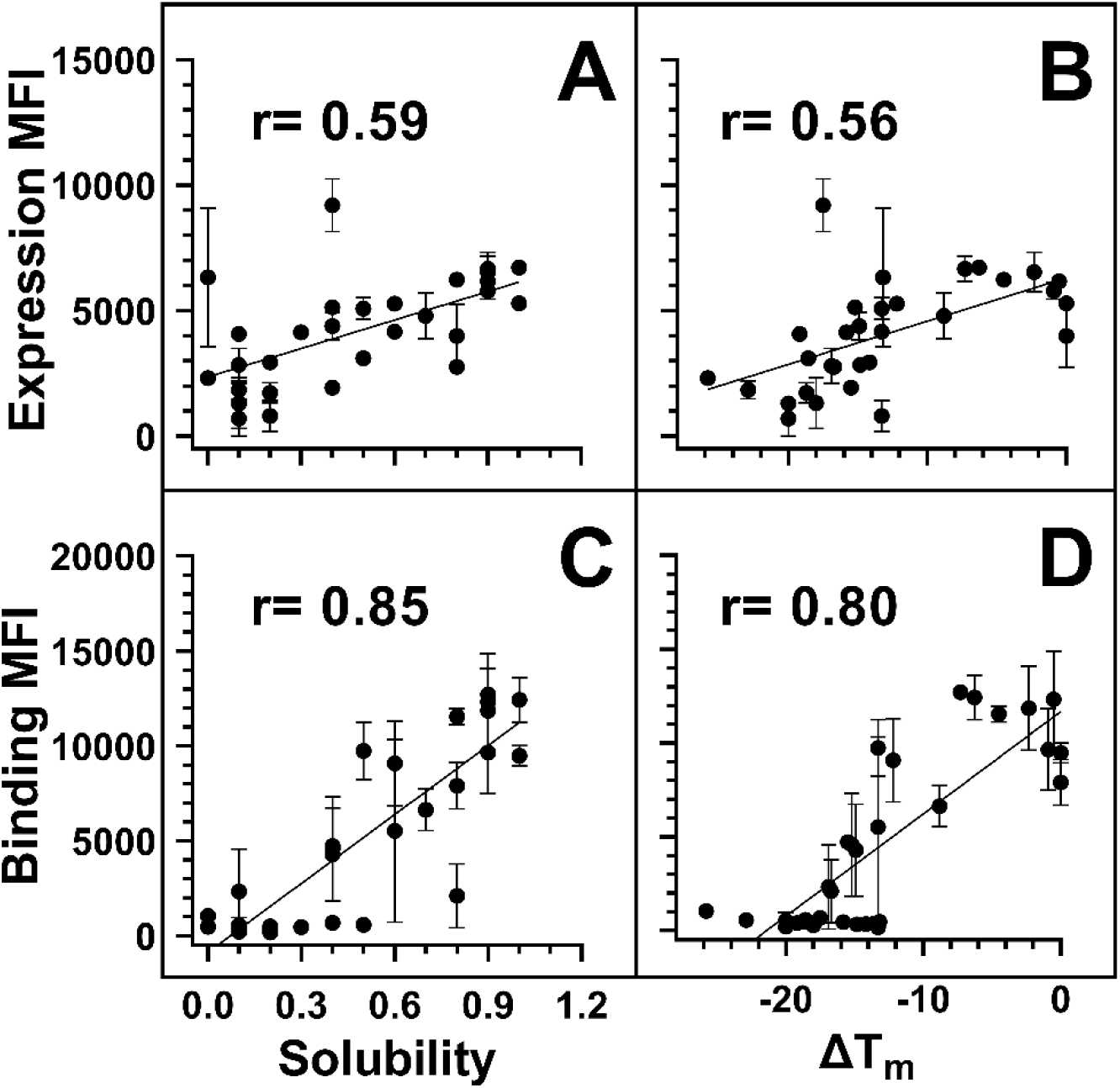
Correlation of *E.coli in vivo* solubility and *in vitro* thermal stability with the amount of total and active protein on the yeast cell surface. For individual mutants, MFI‘s of expression and binding were estimated by probing the HA tag on surface expressed protein and the FLAG tag on cell surface bound GyrA14 respectively. Correlation of the total amount of protein displayed on the yeast cell surface with (A) *in vivo* solubility or (B) ΔT_m_ (T_m_ (mutant)- T_m_ (WT)) of CcdB mutants. Correlation of the amount of active protein on the yeast cell surface with (C) *E.coli in vivo* solubility or (D) ΔT_m_ of CcdB mutants. A better correlation was observed between biophysical parameters with binding MFI rather than expression MFI. In the figure, the ΔT_m_ of WT was increased by 1^○^C to remove overlap with another point. Data for *E.coli in vivo* solubility and thermal stability was taken from Tripathi et al (Tripathi *et al*, 2016). WT data is shown in open circles.

### Deep sequencing analysis of CcdB library and MFI calculation for CcdB mutants

To extend these results, an SSM library of ccdB was expressed on the yeast cell surface. Different populations based on extent of binding to gyrase or cell surface expression were sorted. A total of 32 different populations were sorted at two different concentrations of GyrA14 (100 nM, 5 nM) as a function of either surface expression level or the extent of binding to GyrA14 (Supplementary Figure S2). The lower concentration of GyrA14 was chosen to be around the K_D_ of CcdB-GyrA binding (Supplementary Figure S3), the higher concentration was one where WT CcdB approaches saturation in binding with GyrA14 on the yeast cell surface. We hypothesized that at lower concentrations of GyrA14, the binding on the yeast cell surface will be a function of both stability as well as binding affinity. However, at saturating concentration of GyrA14, the binding on the yeast cell surface will largely be a function of amount of correctly folded protein that in turn might be a function of protein stability, rather than the K_d_ of the mutant(s). MFI was calculated for each mutant as explained in the Methods section. The MFI was calculated at different stringencies (where the stringency refers to the sum of reads for a given mutant over each gate of the histogram), namely 25, 50, 100, 150 and 200 reads. All mutants with a total read number less than the stringency value were removed from the analysis. As the stringency increased, the pairwise correlation between the biological replicates increased (Supplementary Figure S4, Supplementary Table S1). The data was analysed with a stringency of 50 reads, since at higher stringencies, correlation did not improve significantly, but the number of mutants reduced. Reconstructed Binding and Expression MFI from deep sequencing data are hereafter referred to as MFI_seq_ (bind) and MFI_seq_ (expr) respectively.

### MFI reconstruction and its correlation with stability, solubility and residue burial

A few published studies have described estimation of MFI values using deep sequencing of sorted populations and are therefore similar to our experimental strategy. However, the procedure for MFI reconstruction in these reports was relatively complicated compared to that used here (Sharon *et al*, 2012; Peterman & Levine, 2016; Cambray *et al*, 2018; Noderer *et al*, 2014). In those studies, the fractions of reads were calculated in each bin for all the mutants and MFI (mlMFI) of mutants were calculated by fitting the data to a maximum likelihood distribution of the histogram. We found that if mutants are present in only one bin (highly destabilized or nonsense mutants) then this method is unable to perform the MFI calculation (Starr *et al*, 2020). For the remaining mutants we found a good correlation between MFI_seq_ and mlMFI for binding at 5 and 100 nM GyrA14, and for expression (Supplementary Figure S5). For mutants with over 50 reads, we could calculate the MFI of 11,153 mutants using the maximum likelihood method and 11,436 mutants using our method. We also found that progressively reducing the number of bins from eleven to six, does not significantly affect the estimated MFI values, however a further reduction to four bins results in a noticeable change in the estimated values using either method (Supplementary Figure S6). A good correlation was also found between the MFI of individually analysed mutants and their corresponding MFI_seq_ values, validating our approach of MFI reconstruction (Supplementary Figure S7A, 7B). Individually analysed mutants showed a good correlation between the amount of active protein on the cell surface and *in vitro* measured thermal stability of the purified protein. Similarly, we also found a good correlation between MFI_seq_ (bind) of mutants inferred from deep sequencing, and thermal stability as well as *in vivo* solubility for the selected mutants (Supplementary Figure S7C, 7D).

For the exposed residues (>10% accessibility), mutations did not affect the degree of surface expression and binding to GyrA14 (Figure 3A, B). Expression was also unaffected by mutations in the active-site residues (identified from PDB ID:1X75) (Figure 3C). However, many buried site mutants showed very low expression, possibly because of aggregation and degradation inside cells or during export (Figure 3C). In the case of binding for buried and active-site residues, a very high mutational sensitivity was found (Figure 3D) similar to the previous report of CcdB mutants in *E.coli* (Tripathi *et al*, 2016). We also found a very high mutational sensitivity of binding for a few non-interacting residues in the loop connecting beta strands S2 and S3 at both 5 nM and 100 nM GyrA14 concentration (Supplementary Figure S8). The residues I24, I25 and D26 in this loop are directly involved in interacting with Gyrase and mutation at non-interacting residues (22, 23 and 27) in the loop might restrict or alter the conformation of the loop, thus reducing the affinity of CcdB mutants to GyrA14. However, there was no effect on the expression of the mutants in this loop, indicating that the mutant proteins are not destabilized (Supplementary Figure S8). We did not find a high correlation between MFI_seq_ (bind) and either accessibility or depth, because many mutations at both buried and active-site residues have high mutational sensitivity (Supplementary Table S2). The previously described parameter RankScore, is a measure of mutant activity in *E.coli* (Adkar *et al*, 2012) with high RankScore denoting lower activity. We found a poor correlation between the MFI_seq_ (bind) values of CcdB mutants at both exposed non active-site as well as active-site residues, and RankScore. In *E.coli,* most of the exposed non active-site residues do not show any mutational sensitivity, i.e. they have the same RankScore values as WT. However, in the present case many such CcdB mutants show lower binding to GyrA14 compared to WT. The loss of binding could be attributed to the decrease in the affinity between CcdB and Gyrase, or destabilization due to mutation. We defined a new parameter MrMFI (mean residue MFI) which is the mean of the MFI values of all the mutants at a certain position. MrMFI (expr) and MrMFI (bind) at 100 nM GyrA14, show a good correlation with RankScore. (Supplementary Table S2). MrMFI (expr) also showed good correlation with Depth which is a structural measure of residue burial (Chakravarty & Varadarajan, 1999). However, in the case of binding at 5 nM, a weaker correlation of MrMFI (bind) with the aforementioned parameters was observed (Supplementary Table S2). In previous studies, identification of the active-site residues solely from the deep sequencing data was not very efficient (Adkar *et al*, 2012; Bhasin & Varadarajan, 2021), this is presumably because *in vivo* activity is often governed by threshold effects, and because mutations at buried residues also affect activity. The current methodology removes such drawbacks. We could distinguish between buried and active-site residues by comparing the MFI_seq_ (bind) and MFI_seq_ (expr). Most buried site residues showed low values of both MFI_seq_ (bind) and MFI_seq_ (expr) compared to WT. However, the active-site residues showed low MFI_seq_ (bind) but similar MFI_seq_ (expr) compared to WT. We found that the average MFI_seq_ values of charged residues are a good predictor to discriminate between buried and active-site residues. For calculating MrMFI_charged_ of charged WT residues, we only consider mutants with opposite charge. For some mutants at buried positions, we found a very low MrMFI_charged_ (expr) but the mutants were absent in MrMFI_charged_ (bind). We found that such mutants had very high reads, suggesting that the values of MrMFI_charged_ (expr) are correct. We anticipated that such mutants lack binding and are therefore present only in the bin which had a background level of binding signal, the presence of mutant in only that gate led to the removal of such mutants due to the stringency set for the analysis. Hence, such mutants were assigned a MrMFI_charged_ (bind) similar to other buried positions. MrMFI_charged_ had a bimodal distribution (Supplementary Figure S9), so k-means clustering was performed to identify the mean (µ) and standard deviation (σ) of each distribution. The distributions were named D1 (higher MrMFI_charged_) and D2 (lower MrMFI _charged_). Buried site residues were assigned to be those which have MrMFI_charged_ (bind)) and MFI_seq_ (expr) less than the set threshold (µ+0.5*σ) for distribution D2. Active-site residues were assigned as those which had MrMFI_charged_ (bind)) less than (µ+σ) of the D2 distribution and MFI_seq_ (expr) higher than (µ-2*σ) of distribution D1 (Figure 4). The accuracy, specificity and sensitivity of prediction of exposed non active-site, buried and exposed active-site residues are mentioned in Supplementary Table S3.

**Figure 3:**
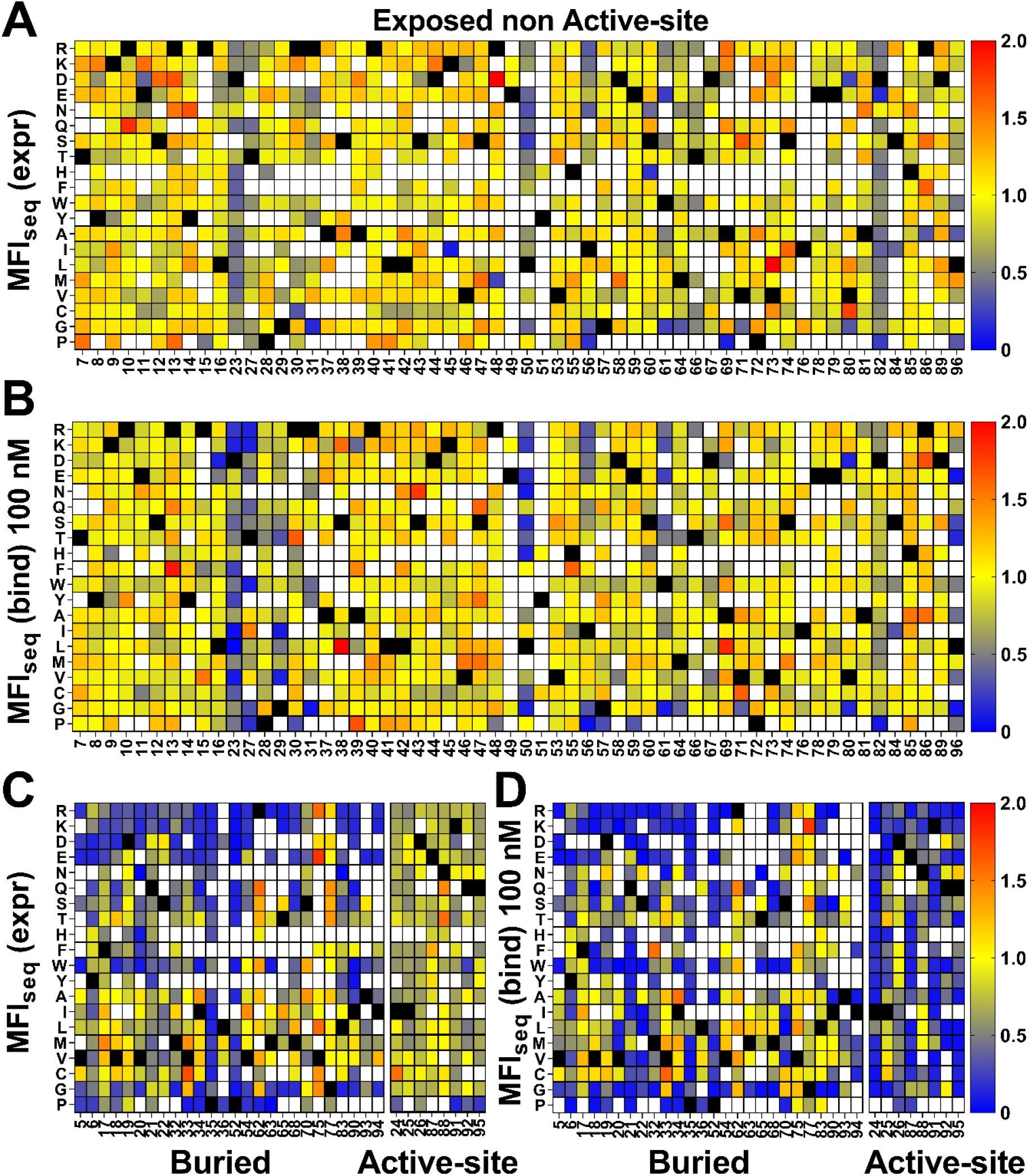
Heatmap of normalized MFI_seq_ values for CcdB mutants. MFI_seq_ value of mutant was divided by the MFI_seq_ value of WT to normalize it. (A) MFI_seq_ (expr) and (B) MFI_seq_ (bind) at 100 nM GyrA14 for exposed non active-site residues. (C) MFI_seq_ (expr) and (D) MFI_seq_ (bind) for buried and active-site residues. Exposed, buried (PDB ID:3VUB) and active-site (PDB ID:1X75) residues are segregated based on the crystal structure. Residues which had accessibility greater than 10% were considered exposed, all remaining residues were considered buried, and active-site mutants in contact with GyrA14 were identified as explained the Methods section. Blue to red colour represents increasing normalized MFI_seq_ values, black colour shows the WT residue at the corresponding position. White colour indicates that the mutant is not available. The buried site residues have very high mutational sensitivity both in case of expression and binding. The active-site residues show mutational sensitivity only with respect to Gyrase binding. Information about the mutational sensitivity of expression and binding can be used to differentiate exposed, buried and active-site residues.

**Figure 4:**
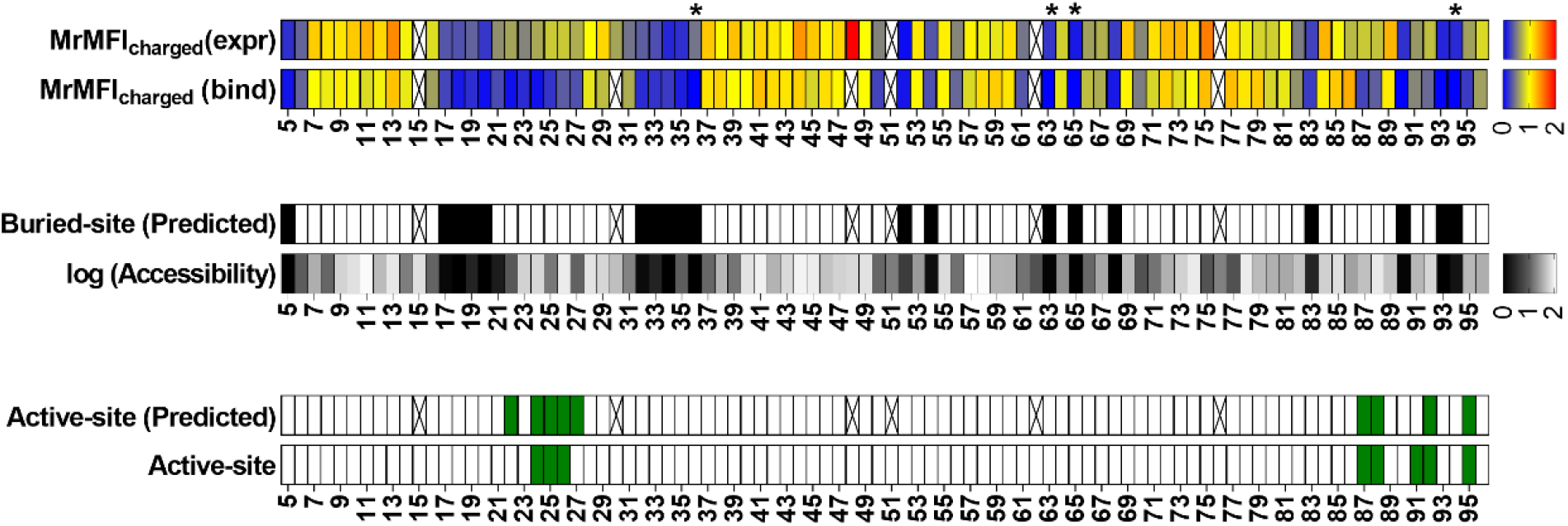
Identification of buried and active-site residues from MrMFI_charged_ (bind) and MrMFI_charged_ (expr). Side chain accessibilities in dimeric CcdB (PDB: 3VUB), darker to lighter shade indicate increasing accessibility, accessibility is reported as log accessibility. the mutants were clustered into two bins based on the distribution of MrMFI_charged_ and k-means and standard deviations were calculated for both distributions. The distributions were named D1 (higher MrMFI_charged_) and D2 (lower MrMFI_charged_). Residues which had MrMFI_charged_ (binding) and MrMFI_charged_ (expr) lower than (µ+0.5*σ) of distribution D2 were characterized as buried. The false negatives were Y6, D19, Q21, S22, S70, V75 and G77, the polar side chains of these residues are pointing towards the surface. Active-site residues were identified as those in contact with GyyrA14 (PDB ID 1X75). Residues which had MrMFI_charged_ (binding) less than (µ+σ) of D2 distribution and MrMFI_charged_ (expr) higher than (µ-2*σ) of distribution D1 were predicted as active-site. We obtained a few putative false positives. However, these residues are likely involved in functional aspects of activity that cannot be inferred from the CcdB:GyrA14 crystal structure. The same residues were seen to be important for CcdB activity *in vivo* in *E.coli* (Tripathi *et al*, 2016). Some positions could not be categorized due to lack of reads, such positions are indicated with an ‘X’. Positions indicated with ‘*’ are the one where MrMFI_charged_ (expr) was observed and the mutants had high read counts but the mutants were absent in MrMFI_charged_ (bind), such positions were assigned MrMFI_charged_ (bind) values similar to other buried positions.

### Selection and characterization of putative stabilized mutants from deep sequencing data

In the previous section, we discussed the correlation between protein biophysical properties such as thermal stability and *in vivo* solubility with either the amount of active protein or the ratio of active protein to total protein on the yeast cell surface for a few (30) mutants. However, most of these mutants were destabilized with respect to the WT protein. To confirm whether this correlation also holds for mutants that have stability similar or greater than WT, we selected a few CcdB mutants based on either the MFI_seq_ (bind) or MFI_seq_ (ratio) (MFI_seq_ (bind)/ MFI_seq_ (expr)) for *in vitro* characterization of thermal stability. We examined the average and standard deviation of expression for all mutants and selected only those mutants based on MFI_seq_ (ratio) which cross a minimum cut-off (µ+0.5*σ) for MFI_seq_ (expr) to remove the bias created by mutants which have very low expression. No threshold for expression was set for selection of mutants based on their MFI_seq_ (bind). No selection of the mutants was performed based solely on the MFI_seq_ (expr).

Six mutants were characterized using the criteria MFI_seq_ (bind) at 5 nM GyrA14, none of them showed a higher T_m_ than WT (Figure 5A); whereas two of the mutants selected on the basis of MFI_seq_ (ratio) showed a significantly higher T_m_ than WT (Figure 5B). A subset of seven mutants was selected based on MFI_seq_ (bind) at 100 nM GyrA14, none of the mutants showed higher stability than WT CcdB (Figure 5C). Ten mutants were selected based on MFI_seq_ (ratio) and characterized, four showed higher stability, two mutants were similar to WT and the remaining four were less stable than WT CcdB (Figure 5D). We therefore hypothesize that if the stability of a mutant crosses a threshold then its expression will not increase further. To confirm this hypothesis, we measured the amount of active protein on the yeast cell surface for seven individual mutants which had T_m_’s ranging from 60 ^ᴼ^C to 70 ^ᴼ^C, and found that the expression and binding for these mutants are similar to each other and to WT (Supplementary Figure S10).

**Figure 5:**
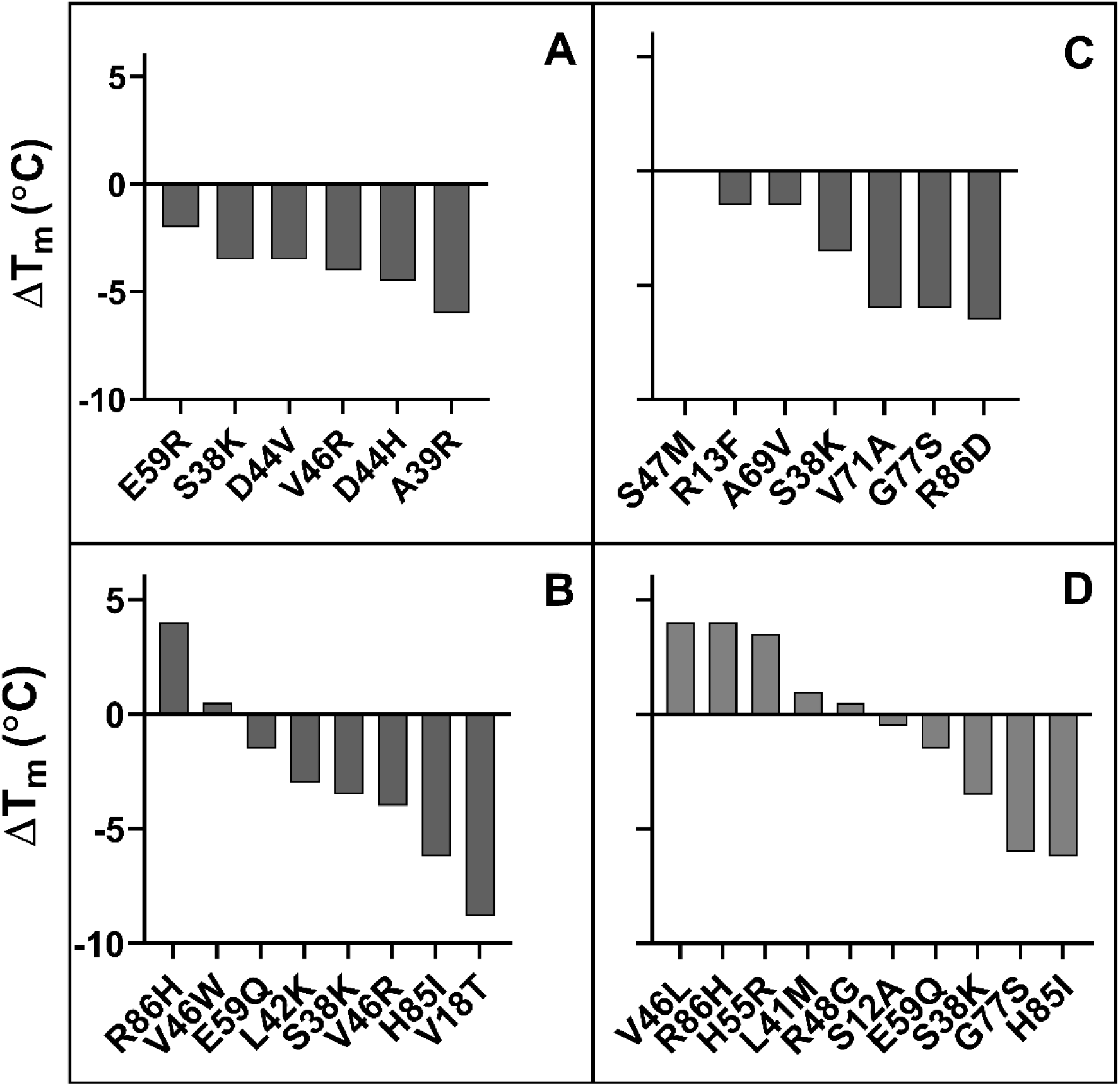
ΔT_m_ of putative stabilized CcdB mutants. Mutants were identified from A) MFI_seq_ (bind) at 5 nM GyrA14, (B) MFI_seq_ (ratio) at 5 nM GyrA14, (C) MFI_seq_ (bind) at 100 nM GyrA14, (D) MFI_seq_ (ratio) at 100 nM GyrA14. The mutants were randomly selected from a subset of forty mutants which showed the highest MFI_seq_ (bind) or the highest MFI_seq_ (ratio) and had MFI_seq_ (expr) > 6672.

### Prediction of thermal stabilities of putative destabilized mutants

For destabilized mutants we observed a good correlation between MFI_seq_ (bind) and T_m_ of individual mutants (Supplementary Figure S7D). Using this correlation, we next predicted the T_m_ of each mutant for an additional set of (n=28) previously described CcdB mutants (Tripathi *et al*, 2016) based on their MFI_seq_ (bind). We found a good correlation (r=0.82) between predicted and *in vitro* measured T_m_ for this set of CcdB mutants as well (Supplementary Figure S11A). This now allows us to identify putative destabilized mutants and accurately predict the extent of destabilization for all such mutants in the CcdB YSD library. We also predicted the thermal stability of CcdB mutants using the *in silico* tool HoTMuSiCv1.0 (Pucci *et al*, 2020), however, we did not find a good correlation between measured and predicted T_m_ (Supplementary Figure S11B). It has been shown that *in vitro* protein thermal stability and free energy of unfolding are correlated (Chen *et al*, 2000; Prajapati *et al*, 2007; Tripathi *et al*, 2016). We therefore predicted the free energy of unfolding for CcdB mutants using SDM (Pandurangan *et al*, 2017), mCSM (Pires *et al*, 2014a), PoPMuSiC (Dehouck *et al*, 2011), DynaMut (Rodrigues *et al*, 2018), DUET (Pires *et al*, 2014b), MAESTROweb (Laimer *et al*, 2016), DeepDDG (Cao *et al*, 2019), CUPSAT (Parthiban *et al*, 2006), PremPS (Chen *et al*, 2020) and INPS-MD (Savojardo *et al*, 2016). We found moderate correlations, with DeepDDG performing the best (r=0.59), but still poorer compared to our prediction from YSD data (r=0.82). For a more detailed comparison we analysed the predictions of stability by DeepDDG, since this showed the highest correlation with measured stability of individual mutants at non active-site residues. We excluded residues 21, 22, 23 and 27 as these positions behaved like active-site residues. We found that trends for ΔΔG predicted by DeepDDG for exposed non active-site residues are similar to those obtained from MFI_seq_ (bind) (Figure 6A, 6B). However, we observed some mutant specific differences at residues 8, 16, 50, 53 and 96. Mutations at residues 50 and 96 have highly deleterious effects which reduced GyrA14 binding to yeast surface displayed protein, these are only partially predicted by DeepDDG. In the case of charged and polar mutations at residue 8, 16 and 53 we did not observe a reduction in binding, but the software predicted them to be destabilizing. In the case of buried positions, we found mutation specific effects at 35, 52 and 94 where DeepDDG predicted changes were significantly smaller than the experimentally observed ones. We also found that most of the phenylalanine, tryptophan and arginine mutations were highly destabilizing and the mutants did not bind to GyrA14, however the software gave a lower stability penalty for these substitutions (Figure 6C, 6D). Our MFI based measurements suggested greater destabilization for several mutants relative to DeepDDG prediction. While the overall trends were similar, as discussed above, there are several differences between MFI based and DeepDDG based stability predictions.

**Figure 6:**
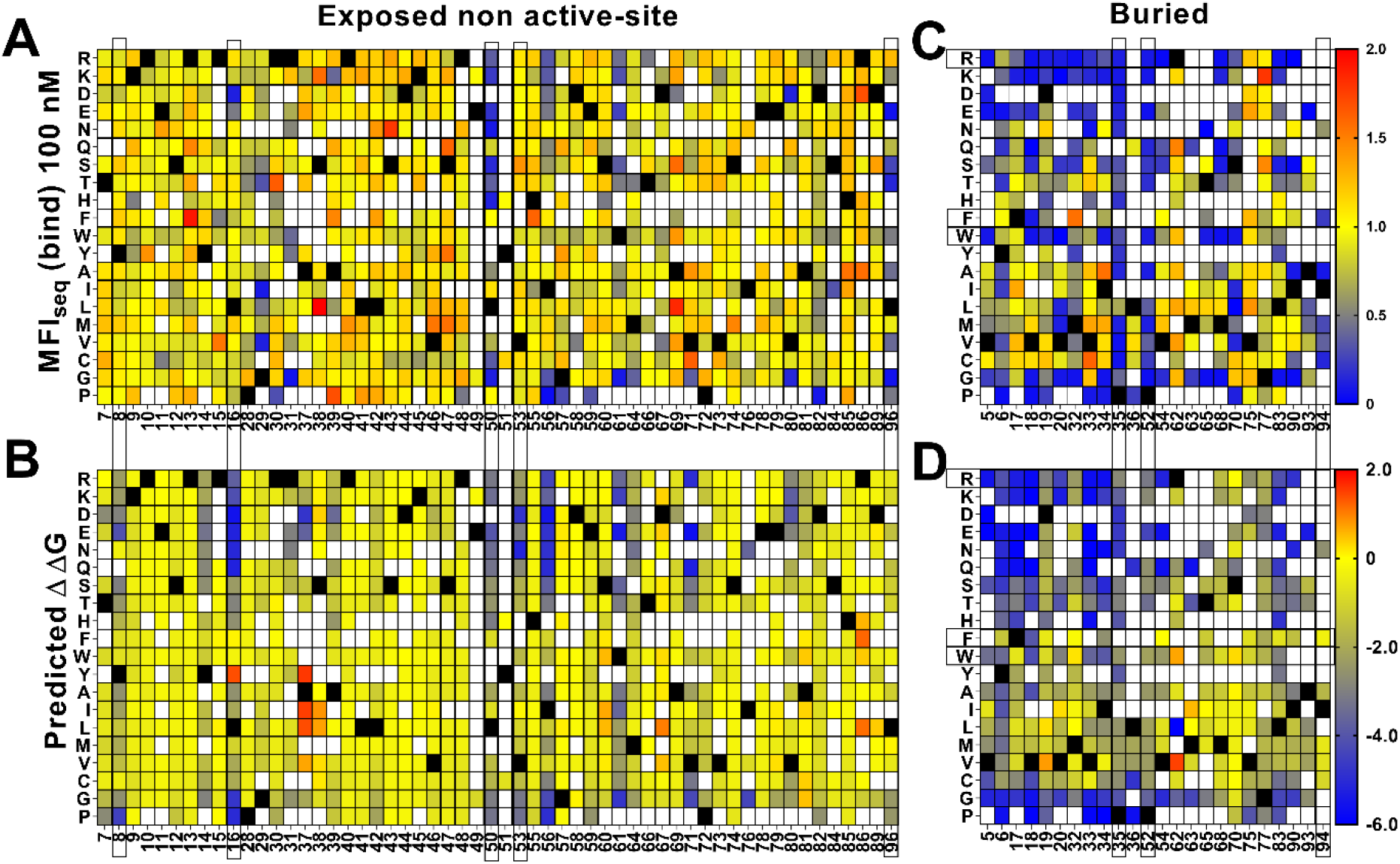
Comparison of stabilities estimated by DeepDDG and yeast surface display. Heat maps for (A, C) MFI_seq_ (bind) normalized to WT and (B, D) ΔΔG predicted by DeepDDG. Residue positions or specific amino acid mutations showing significantly different predicted stabilities by the two methods are highlighted by a box. Blue to red colour corresponds to increasing stability.

### Deep mutational scanning of SARS COV-2 receptor binding domain (RBD)

To examine the generality of our approach, we also analyzed recently reported deep mutational scanning data of the SARS-CoV-2 receptor binding domain (Starr *et al*, 2020). In this study two separate libraries were generated and individually sorted based on expression and binding to ACE-2. The binding (Sortseq (bind)) or expression ((Sortseq (expr)) MFIs relative to WT for barcoded mutants were calculated from the deposited NGS data as explained in the Methods section. Additionally, we analyzed binding at only one concentration of ACE-2 (100 pM, TiteSeq_09) at which the binding started to saturate. Buried residues were those with <10% side chain accessibility in chain C of PDB ID 7KMH (Jones *et al*, 2020a). ACE-2 binding (active-site) residues were assigned as those contacting ACE-2 (Malladi *et al*, 2021). To identify the active-site and buried residues from Sortseq data, we calculated the MrMFI_charged_ for each position. Similar to CcdB, we observed a bimodal distribution for both MrMFI_charged_ (bind) and MrMFI_charged_ (expr) (Supplementary Figure S12) and k-means and standard deviation were calculated for both the distribution D1 (higher MrMFI_charged_) and D2 (lower MrMFI_charged_). As described above for CcdB, buried residues were identified as those which had MrMFI_charged_ (bind) and MrMFI_charged_ (expr) less than the set threshold (µ+0.5*σ) for distribution D2. The active-site positions were identified as those which had MrMFI_charged_ (bind) lower than the set threshold (µ+σ) for population D2 and MrMFI_charged_ (expr) values higher then (µ-2*σ) for population D1. We accurately identified most of the buried residues, however there were some false positive and false negative predictions relative to the crystal structure information (Figure 7). We found 21 positions to be false negative buried positions. We categorized these false negatives into two categories, namely, glycine and the side chains which are pointing towards the surface. The accessibility calculated by DEPTH server for glycine was zero and we therefore expected glycine to fall into the false negative buried category. Thirteen positions out of twenty-one false negative were glycine. Another six positions, 336, 348, 361, 443 and 480 had their side chains pointing towards the protein surface. We also found similar false negative buried residues in CcdB where the side chain hydrophilic group was pointing towards the protein surface. Position 363 and 365 in RBD had accessibility <10% and were pointing towards the core of the protein in the PDB (7KMH) used to calculate accessibility. However, we found that these positions have high accessibility (>30%) in another structure (PDB ID 7D2Z). All the available RBD structures are in complex with other molecules, this might be responsible for variation in the accessibility of residues in different RBD structures. We found 17 false positive buried residue predictions, seven of them were aromatic, seven are charged or polar, two are prolines and one is an aliphatic residue. These positions have both reduced expression and binding for charged residue substitutions (Supplementary Figure S13A, 13D) similar to the buried residues (Supplementary Figure S13B, 13E). The specificity, sensitivity and accuracy of prediction is mentioned in Supplementary Table S3. Active site residues were identified with very high accuracy (Supplementary Table S3), though there were a few false negative and false positive predictions. Additionally, we found several positions which had Sortseq (expr) like WT, however, they had very low Sortseq (bind) (Supplementary Figure S13A, 13D). We hypothesize that these positions are also assisting in the maintenance of proper RBM conformation and enabling its binding to ACE-2. Residues 447, 448, 473 and 476 which gave false positive results, 447 and 476 are part of the receptor binding motif (RBM) and contain glycine in a conformation which is available only for glycine. Hence mutation to a non-Gly residue will likely disrupt the conformation of the RBM thus decreasing binding to ACE-2. Mutations at positions 446, 453, 493 and 498 gave false negative results. Of these false negative positions, 446 is again glycine. We found that the Arg mutants at N493 and N498 positions have very little effect on expression and binding (Supplementary Figure S13C, 13F). We hypothesized that these positions may not have the most optimal WT residue, or they may show no mutational penalty for binding to ACE-2. A recent report showed that the affinity of Q498R to ACE-2 is higher than WT RBD (Xue *et al*, 2020) and was enriched as double mutant Q498R/N501Y when selection was performed for RBD mutants having high affinity towards ACE-2 (Zahradník *et al*, 2021). It has also been reported that when chimeric virus evolved in the presence of neutralizing antibodies C121 and C141, this enriched for the Q493R mutation. The mutant virus grows to high PFU titers similar to WT, and infectivity is also inhibited by a chimeric ACE-2 analog, similar to WT (Weisblum *et al*, 2020). The specificity, sensitivity and accuracy of prediction is mentioned in Supplementary Table S3.

**Figure 7:**
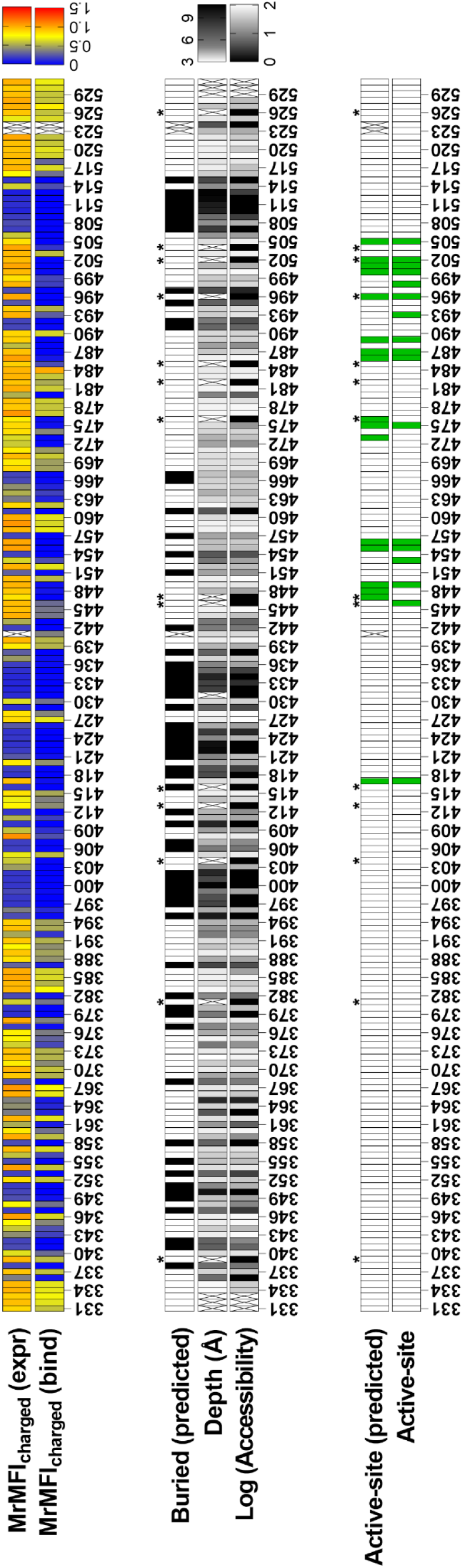
Prediction of buried and active-site positions in SARS-CoV-2 RBD from Sortseq data. Buried residues were identified from chain C of PDB ID 7KMH, residues which had <10% side chain accessibility were categorized as buried. The accessibility and depth was calculated using DEPTH server (Tan *et al*, 2011). Active-site residues were identified from PDB ID 6M0J as explained earlier (Malladi *et al*, 2021). Criteria used to predict buried and active-site positions from MFI data were identical to those used for CcdB. Positions which did not have MrMFI data or could not be assigned to either buried or active-site categories are highlighted with “X”. Accessibility calculated by DEPTH server for glycine is zero and these are marked with a ‘*’.

## Discussion

With the advancement of mutagenesis and directed evolution methodologies, proteins with modified traits and function can be developed in a relatively short duration of time (Bornscheuer *et al*, 2019; Chen & Arnold, 1991; Winter *et al*, 1994). *E.coli* remains an expression host of choice for many proteins and high level, soluble *E.coli* expression is a desirable attribute. When eukaryotic or unstable prokaryotic proteins are overexpressed in bacteria, they often tend to form insoluble aggregates called inclusion bodies (IB). Formation of IBs often results in low yields of purified soluble protein. Designing improved variants of a protein by increasing half-life, stability and activity is an ongoing requirement of most pharmaceutical and biotechnology industries. However, a reliable, high-throughput, efficient and rapid method is required for solubility and stability analysis of engineered proteins. Previously, several high-throughput methods to select for soluble expression have been developed based on fusion to a reporter protein. These rely on the reporter activity, which is perturbed if an aggregation prone protein is fused (Maxwell *et al*, 1999; Wigley *et al*, 2001; Fisher, 2006; Waldo *et al*, 1999). These methods can be used to isolate protein variants with enhanced solubility but cannot reveal if the fused protein is properly folded. In some cases, such unstable proteins may also form soluble aggregates (Tripathi *et al*, 2016). Since many of these reporter screens employ cytoplasmic expression and use bacterial hosts, disulphide rich or glycosylated proteins, or those binding to complex ligands cannot be studied. Yeast surface display coupled to FACS, has been widely used to evolve such targets. Typically, populations are sorted for multiple rounds to enrich for stable binders to a target of interest (Traxlmayr & Obinger, 2012; Esteban & Zhao, 2004; Kim *et al*, 2006; Kieke *et al*, 1999). While this approach readily selects for high affinity binders, selecting for stable proteins is more difficult. In some cases, this methodology has also been used to isolate stable variants of proteins (Pepper *et al*, 2008) and a good correlation was observed between surface expression and improved biophysical parameters. However, other studies in different systems did not find such a correlation (Park *et al*, 2006a; Piatesi *et al*, 2006).

In the present work we utilize YSD to measure the amount of total protein as well as total active protein displayed on the yeast cell surface. A good correlation was found between the amount of active CcdB mutant on the yeast surface and corresponding *in vivo* solubility in *E.coli* (r=0.85) or T_m_ (r=0.80). A recent report also suggests that the amount of active protein on the yeast cell surface can be used as a criterion to isolate stable mutants (Traxlmayr & Shusta, 2017). In the present study, no correlation was found between the amount of total protein on the yeast cell surface and the biophysical properties of mutants. A few mutants which have very low solubility in *E.coli* showed very high expression, but there was a negligible amount of active protein on the yeast surface. It has been previously suggested that the quality control system in yeast is not able to discriminate these mutants from properly folded ones or alternatively that the folded conformation is maintained by chaperones in the ER (Park *et al*, 2006b). Once these mutants are exported to the cell surface they may start to unfold. This could be one reason why some groups including ours did not find a good correlation of surface expression with the stability or solubility of these proteins. In previous studies (Shusta *et al*, 1999), a very limited number of proteins were used for surface expression studies, it is possible that in this small number, mutants which had high surface expression or secretion but lower stability than WT were not observed.

Yeast surface display coupled to FACS typically requires multiple rounds of sorting to enrich variants with desired activity and phenotype. Here, we have performed a single round of sorting and developed a rapid, uncomplicated procedure of estimating MFI’s of individual mutants of CcdB combining FACS and deep sequencing. This MFI_seq_ was shown to correlate well with the corresponding experimentally measured MFIs for several individual mutants. The MFI_seq_ was used to generate the mutational landscape of expression and binding of a mutant library.

We showed that such data can be used to accurately discriminate between buried, exposed non active-site and exposed active-site residues both for CcdB and an unrelated protein, RBD of the spike protein of SARS-CoV-2. Highly destabilizing charged mutations in the core of the protein decreased both expression and binding, while the active-site residues showed reduction in binding alone for charged mutations. Relative to an earlier study which assayed *in vivo* activity in *E.coli* (Adkar *et al*, 2012), the present methodology is better able to identify and distinguish between the two categories of mutationally sensitive residues, namely buried and exposed, active-site residues. Identification of active-site residues of interacting partners through charged mutation scanning provides a better alternative to alanine and cysteine scanning mutagenesis. In general, mutations that affect total activity *in vivo* can do so by affecting specific activity without changing the amount of folded protein, decrease the amount of folded protein without affecting specific activity or a combination of the above. The present analysis distinguishes between the above possibilities, and is therefore able to distinguish buried from exposed, active-site positions. This is useful for applications that attempt to use saturation mutagenesis data for protein model discrimination and structure prediction (Khare *et al*, 2019; Jones *et al*, 2020b) as well as interpreting clinical data on disease causing mutations (Livesey & Marsh, 2020; Findlay *et al*, 2018).

MFI_seq_ (bind) was also used to predict the T_m_ of CcdB mutants. We found a good correlation between predicted and measured ΔT_m_ for a subset of CcdB mutants. We also compared the accuracy of *in silico* tools used to predict the stability of mutants and found that these tools had lower accuracy relative to our approach. We used experimental stability measurements for a small number of destabilized mutations, combined with MFI_seq_ measurement to predict stabilities of all destabilized mutants in the saturation mutagenesis library. We could readily identify destabilized mutants of CcdB, however, the recovery of mutants more stable than WT was lower, but still significant, considering the rarity of such mutations. This is likely due to the possibility that if the stability of the protein crosses a threshold, additional increments in stability do not result in enhanced expression or binding.

A limitation of the present approach is that it requires an epitope tagged or fluorescently labelled conformation specific binding partner. Another limitation could be differential relative stability of proteins upon yeast cell surface display compared to expression in the native host and/or intracellular expression. For glycosylated proteins, the stability of mutants may also be altered because of hyper glycosylation of protein on the yeast cell surface compared to proteins expressed in mammalian systems or prokaryotic systems where glycosylation is absent. The presence of glycosylation may also affect the binding to a cognate partner which in turn may give rise to false results. This does not appear to be the case for the SARS-CoV-2 RBD which contains a single glycan at residue 343, but may be an issue for protein with multiple glycosylation sites. We are examining these possibilities in ongoing studies. Despite these caveats, the present study suggests that the proposed methodology can accurately distinguish buried from active-site residues, quantitatively estimate thermal stabilities of destabilized mutants in large libraries, and also be used with moderate accuracy to identify stabilized mutants.

## Materials and Methods

### Bacterial strains, yeast strains and plasmids

*E.coli* CSH501 strain carries a mutation in the gyrA gene which abolishes inhibition and poisoning by CcdB (Bajaj *et al*, 2008b). The EBY100 strain of Saccharomyces cerevisiae has the aga1 gene under the Gal1 promoter for inducible expression and a TRP1 auxotrophic mutation. The strain lacks the aga2 gene, so only Aga2p fused protein expressed from the plasmid, will form a complex with the Aga1p for yeast cell surface display (Boder & Wittrup, 2000). The ccdB gene was cloned in the pBAD24 plasmid for controllable expression in *E.coli*. ccdB mutants were cloned in the pPNLS shuttle vector for yeast cell surface expression (Najar *et al*, 2017).

### Cloning of WT and mutant ccdB in *E.coli*

ccdB mutants in pBAD24 were generated using three fragment Gibson assembly. Briefly, ccdB was amplified in two fragments using two sets of oligos. For each fragment one of the oligos binds to the vector and the other binds to the gene. The primer of both fragments which bind to the gene were completely overlapping and contained the desired mutation. The fragments were gel extracted and Gibson assembled with NdeI and HindIII digested pBAD24 vector. The Gibson assembled product was electroporated in *E.coli* CSH501 strain and positive transformants were selected on LB agar media containing ampicillin (100 μg/mL). The sequence was confirmed by Sanger sequencing. Sequence confirmed WT or mutant ccdB in pBAD24 vector was used as a template for PCR to amplify the ccdB gene by Vent DNA polymerase. The PCR amplified product was co-transformed with SfiI digested pPNLS vector in the EBY100 strain of S*accharomyces cerevisiae* using LiAc/SS carrier DNA/PEG method for *in vivo* recombination (Gietz & Schiestl, 2007). Positive transformants were selected on SDCAA Tryptophan dropout media plates and the sequence was confirmed by Sanger sequencing.

### Protein Purification

WT and mutant CcdB was purified as described previously (Chattopadhyay & Varadarajan, 2019). Briefly, an overnight culture was diluted 100-fold in LB media containing ampicillin (100µg/ml) and induced with L-arabinose (0.2% w/v) at an OD_600_ of ∼0.5. Following induction for 3 hours, cells were harvested and lysed by sonication. The soluble fraction was separated using centrifugation and incubated with CcdA peptide (residues 45-72^n^) coupled to Affigel-15 at 4 ^ᴼ^C. The unbound fraction was removed and the column was washed with bicarbonate buffer (50 mM NaHCO_3_, 500 mM NaCl, pH 8.5). The bound protein was eluted with 200 mM glycine (pH 2.5) and collected in an equal volume of 400 mM HEPES buffer (pH 8) to neutralize the acidity of glycine.

GyrA14 was purified as described previously (Dao-Thi *et al*, 2004). Briefly, an overnight culture was diluted 100-fold in LB media containing ampicillin (100µg/ml) and induced with IPTG (1 mM) at an OD_600_ of ∼0.5. Following induction for 3 hours, cells were harvested and resuspended in TES buffer (0.2 M Tris, pH 7.5, 0.5 mM EDTA, 0.5 M sucrose and 1 mM PMSF). Cells were lysed and the soluble fraction was separated using centrifugation. The soluble fraction was incubated with pre-equilibrated Ni-NTA beads for 2 hours at 4 ^ᴼ^C. The unbound fraction was removed, and the column was washed with 100 column volumes of wash buffer (50 mM imidazole in 0.05 M Tris, pH 8, 0.5 M NaCl). The protein was eluted with 500 mM imidazole in 0.05 M Tris, pH 8, 0.5 M NaCl and dialysed against 1x PBS.

### Estimation of solubility of WT and mutant CcdB in *E.coli*

*E.coli* CSH501 strain, transformed with pBAD24 plasmid containing WT or mutant ccdB, was grown in media containing ampicillin for 16 hours at 37 ^ᴼ^C and 180 RPM. A secondary culture was grown by diluting overnight grown culture 100-fold. Upon reaching an OD_600_ of 0.4-0.5, CcdB variants were induced with Arabinose at a final concentration of 0.2%(w/v) for 3 hours. The cells were harvested from 1.5 ml culture and lysed in 500µL 1X PBS, using sonication. Supernatant and pellet fractions were separated by centrifugation at 13000 RPM at 4 ^ᴼ^C. The pellet fraction was resuspended in 500 µL 1X PBS and equal volumes of pellet and supernatant fractions were loaded on Tricine-SDS-PAGE to measure the relative amounts of protein in each fraction.

### Protein thermal stability measurement using Thermal shift assay (TSA)

The thermal shift assay was conducted in an iCycle iQ5 Real Time Detection System (Bio-Rad, Hercules, CA). A solution of total volume 20 μL containing 10 μM of the purified CcdB protein and 2.5X Sypro orange dye in suitable buffer (200 mM HEPES, 100 mM glycine), pH 7.5 was added to a well of a 96-well iCycler iQ PCR plate. The plate was heated from 15 ^ᴼ^C to 90 ^ᴼ^C with a 0.5 ^ᴼ^C increment every 30 seconds. The normalized fluorescence data was plotted against temperature and T_m_ measured as described (Niesen *et al*, 2007; Tripathi *et al*, 2016).

### Yeast surface expression of WT and mutant CcdB proteins in EBY100 cells and flow cytometric analysis

*Saccharomyces cerevisiae* EBY100 cells containing WT ccdB or mutant in pPNLS plasmids were grown in three ml SDCAA media (glucose 20g/L, yeast nitrogen base 6.7g/L, casamino acid 5g/L, citrate 4.3g/L, sodium citrate dihydrate 14.3g/L) for sixteen hours. Grown cells were diluted to an OD_600_ of 0.2 in three ml SDCAA media and grown till the OD_600_ reached two. Thirty million cells were harvested using centrifugation and resuspended in three ml SGCAA induction media (galactose 20g/L, yeast nitrogen base 6.7g/L, casamino acid 5g/L, citrate 4.3g/L, sodium citrate dihydrate 14.3g/L) for sixteen hours at 30 ^ᴼ^C, 250 RPM (Chao *et al*, 2006). One million cells were used for flow cytometric analysis. The amount of total protein expressed on the yeast cell surface was estimated by incubating the induced cells in 20 μL FACS buffer (1X PBS and 0.5% BSA), containing chicken anti-HA antibodies from Bethyl labs (1˸600 dilution) for 30 minutes at 4 ^ᴼ^C. This was followed by washing the cells twice with 100 μL FACS buffer at 4 ^ᴼ^C. Washed cells were incubated with 20 µL FACS buffer containing goat anti-chicken antibodies conjugated to Alexa Fluor 488 (1:300 dilution), for 20 minutes at 4 ^ᴼ^C. Fluorescence of yeast cells was measured by flow-cytometric analysis. The total amount of active protein on the yeast cell surface was estimated by incubating the induced cells in 20 μL FACS buffer containing 100 nM GyrA14 for 45 minutes at 4 ^ᴼ^C. Cells were washed and incubated with 20µL mouse anti-FLAG antibodies (1˸300). This was followed by washing the cells twice with FACS buffer, followed by incubating with 20 µL rabbit anti-mouse antibodies conjugated to Alexa Fluor 633 (1:1600 dilution). The flow-cytometric analysis was carried out on BD Accuri or BD Aria III instruments.

### Yeast surface expression and sorting of CcdB Single-site saturation mutagenesis (SSM) library

Previously, an SSM library of ccdB was generated in the pBAD24 vector (Adkar *et al*, 2012; Tripathi *et al*, 2016). The library was PCR amplified using primers having homology to the pPNLS vector. The PCR amplified library was gel extracted and cloned in pPNLS vector using yeast *in vivo* recombination.

A similar protocol was used for sample preparation of the library for FACS as described above for the single mutants with slight modifications. Briefly, ten million cells were taken for FACS sample preparation and the reagents were used in 10X higher volumes compared to the earlier flowcytometric analysis. Two different concentrations of GyrA14 (100 nM, 5 nM) were used for sorting CcdB mutants based on the binding in the 1D histogram. The cells were sorted in 11 and 10 different populations (bins) in case of binding with GyrA14 at concentrations of 100 nM and 5 nM respectively. Additionally, 11 different populations (bins) were sorted from the expression histogram. The experiment was repeated in a biological replicate. The sorting of CcdB libraries was performed using a BD Aria III cell sorter.

### Sample preparation for deep sequencing

Sorted populations were grown on SDCAA agar plates for 48 hours. Colonies were scraped and plasmids were extracted from the cells. The ccdB gene was PCR amplified using primers which bind upstream and downstream of the ccdB sequence and had multiplex identifier (MID) sequence to segregate the reads from different sorted bins. The DNA was amplified for 15 cycles using PCR and the amplified product was gel extracted and purified. Equal amounts of DNA from each sorted population were pooled, and the library was generated using the TruSeq™ DNA PCR-Free kit from Illumina. The sequencing was done on an Illumina HiSeq 2500 250PE platform at Macrogen, South Korea after incorporating 20% ϕX174 DNA in the library.

### Analysis of deep sequencing data

Deep sequencing data for the ccdB mutants obtained from the Hiseq 2500 platform was processed using a pipeline developed by adopting certain aspects from an already existing in-house protocol (https://github.com/skshrutikhare/cys_library_analysis). The latter method involved the alignment with wild type sequence followed by merging of the paired-end reads, while in the modified protocol, the reads are first merged and then aligned with the wild-type sequence. The present methodology consists of the following steps: assembling the paired end reads, quality filtering, binning, alignment and mutant identification. All these steps were incorporated in a pipeline and made executable from a single command using a parameter file unique to a given data-set. In the first step, paired end reads were assembled using the PEAR v0.9.6 (Paired-End Read Merger) tool (Zhang *et al*, 2014). The “quality filtering” step involved deletion of terminal “NNN” residues in the reads, and removal of reads, not containing the relevant MID and/or primers, along with the reads having mismatched MID’s. Finally, only those reads having bases with Phred score ≥ 20 are retained. A binning step involved further filtering, which eliminated all those reads having incorrectly placed primers, truncated MIDs/primers (due to quality filtering) and shorter/longer sequences than the length of the wild type sequences. The remaining reads were binned according to the respective MIDs. In the alignment step, reads were aligned with the wild type ccdB sequence using the Water v6.4.0.0 program (Smith & Waterman, 1981) and reformatted. The default values of all parameters, except the gap opening penalty, which was changed to 20, was used. In the final step of “substitution”, reads were classified based on insertions, deletions and substitutions (single, double etc mutants).

### MFI reconstruction from deep sequencing data

Reads of each mutant were normalized across different bins individually (Equation 1), and the fraction of each mutant (*Xi*) distributed amongst the different bins was calculated (Equation 2). The reconstructed MFI for an individual mutant was calculated by the summation of the product, obtained upon multiplying the fraction (*Xi*) of the mutant in a particular bin (*i*) with the MFI of the corresponding bin obtained from the FACS experiment (*Fi*), across the various bins populated by the respective mutant (Equation 3).

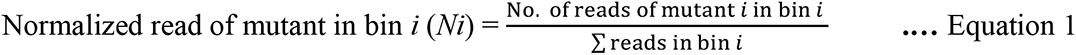

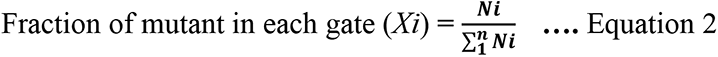

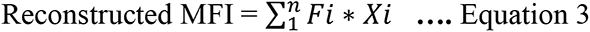

The MFI_seq_ of the biological replicates were different so the MFI_seq_ of one of the replicates was adjusted using “m” and “c” obtained from the correlation between the replicates and then averaged.

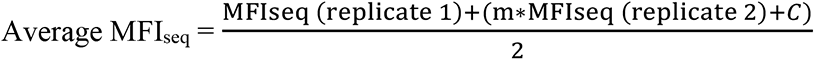

### Maximum likelihood MFI (mlMFI) calculation

Reads of each mutant were normalized within and across the bins. The fraction of each mutant (*Xi*), distributed amongst the different bins, was calculated as explained in the above section. The fraction (*Xi*) was multiplied with a scaling factor to convert the data into integers as this is required by the package below. The mlMFI was calculated using a maximum likelihood method using the fitdistrplus R package as explained earlier (Starr *et al*, 2020). The ‘fitdistcens’ function in the fitdistplus R package helps in the estimation of fluorescence values for such observations using a maximum likelihood approach, where the values are transformed into a data frame of two columns left and right, describing each observed value as an interval and assuming a normal distribution of values. The left column contains the left bound of the interval and the right column contains the right bound of the interval for interval-censored observations, based on the fluorescence boundaries of each bin. The maximum likelihood approach was used to estimate the MFI of binding and expression for each mutant, based on its distribution of reads across the sorted bins, and the fluorescence boundaries of each sorted bin.

### MFI calculations after bins merging

The bins were merged following which mlMFI amd MFI_seq_ were calculated for GyrA14 binding (100 nM) for replicate 1. The fraction of each mutant in each bin was calculated as explained in the earlier sections. To merge bins for a given mutant, fractions present in each of the bins to be merged were added arithmetically. For mlMFA calculation, the minimum and maximum fluorescent boundary of the merged bin was set at the lowest and highest value of the fluorescent boundary for that set of bins. The mlMFI of CcdB mutants was calculated as explained above. In the case of MFI_seq_, the mean fluorescent intensity of merged bins was determined by making a new bin spanning the set of merged bins. The MFI_seq_ of CcdB mutants was then calculated as explained above.

### Depth, accessibility and RankScore calculations

Depth was calculated using the server DEPTH (Tan *et al*, 2011; Chakravarty & Varadarajan, 1999). Accessibility was calculated using the program NACCESS (Hubbard SJ, 1993). In both cases, the input co-ordinates were homodimeric CcdB (PDB ID 3VUB). RankScore and MS_seq_ are measures of mutational sensitivity in *E.coli*. Values were obtained from Adkar et al (Adkar *et al*, 2012). Buried residues were those with <10% accessibility in 3VUB. Active-site residues were those with ΔASA>0. ΔASA difference between the solvent accessible surface area of CcdB residues in the free (3VUB) and GyrA14-bound forms (1X75) respectively (Aghera *et al*, 2020).

### Deep mutational scanning of SARS COV-2 receptor binding domain (RBD)

The deep mutational scanning data was taken from a recent report (Starr *et al*, 2020) in which two independent libraries of RBD were generated and sorted in four different bins based on expression or binding to ACE-2. In the MFI of binding and expression for individual mutants was reconstructed in that study using a maximum likelihood method using fitdistrplus R package. The expression MFI (Sortseq (expr)) data was shared by the authors in a repository (https://github.com/jbloomlab/SARS-CoV-2-RBD_DMS). We reconstructed the binding MFI (Sortseq (bind)) at an ACE-2 concentration of 100 pM (TiteSeq_09). For Sortseq (bind) estimation we used the script provided by the authors (https://github.com/jbloomlab/SARS-CoV-2-RBD_DMS/blob/master/results/summary/compute_expression_meanF.md). The authors used data from both single and multiple mutants, together with a model to account for epistatic effects to infer the MFI values for individual mutants. We modified the script to change the input data required to calculate Sortseq (bind). For both Sortseq (bind) and Sortseq (expr), we analyzed only single mutant data to avoid any artifacts that might arise from the epistatic model and took the average of delta Sortseq MFI (log(Sortseq (WT)) – log(Sortseq (mutant))) of mutants which had multiple barcodes. The Sortseq MFI values of mutants were averaged between the two libraries and the antilog was calculated for delta Sortseq MFI to analyse the ratio of Sortseq (bind) or Sortseq (expr) of mutants with respect to WT.

## Data availability statement

The deep sequencing data discussed in the present study has been deposited in NCBI’s Sequence Read Archive (accession no. SRR16071134). Illumina sequencing counts for each ccdB mutant of FACS bins are available at https://github.com/rvaradarajanlab/ccdb_ssm/blob/main/ccdb_SSM_lib_reads.xlsx. MFI_seq_ and mlMFI of CcdB mutants are available at https://github.com/rvaradarajanlab/ccdb_ssm/tree/main/Calculated_MFI. FCS files of individual mutant expression and binding estimated using flow cytometry is available at https://github.com/rvaradarajanlab/ccdb_ssm/tree/main/Individual_mutant_exp_binding. FCS files of GyrA14 titration against WT CcdB on the yeast cell surface using flow cytometry are available at https://github.com/rvaradarajanlab/ccdb_ssm/tree/main/CcdB-GyrA14_Kd.

Codes used to calculate expression and binding mlMFI using maximum likelihood method is available at https://github.com/rvaradarajanlab/ccdb_ssm/tree/main/scripts. the remaining data is available in the manuscript.

### Supporting information

This article contains supporting information.

## Acknowledgements

S.A. acknowledges Council of Scientific & Industrial Research for his fellowship (SPM-07/079(0218)/2015-EMR-I). K.M. is thankful to Department of Science and Technology (DST) Science and Engineering Research Board for financial support, sanction order no: PDF/2017/002641. M. B. acknowledges Council of Scientific & Industrial Research for her fellowship (SRF-09/079(2766)/2017-EMR-I). Aparna Asok is duly acknowledged for FACS.

## Author contribution

R.V. and S.A. designed the experiments. S.A. performed all the experiments, R.V. and S.A. analyzed all the data. K.M. wrote the software and carried out the processing of the deep sequencing data. M. B. calculated the MFI of CcdB mutants using maximum likelihood method. R.V. and S.A. wrote most of the manuscript.

## Funding and additional information

This work was funded by grants to RV from the Department of Science and Technology, grant number-EMR/2017/004054, DT.15/12/2018), Government of India, Department of Biotechnology, grant no. BT/COE/34/SP15219/2015 DT. 20/11/2015, Ministry of Science and Technology, Government of India and Bill and Melinda Gates Foundation (USA) (INV-005948). We also acknowledge funding for infrastructural support from the following programs of the Government of India: DST FIST, UGC Centre for Advanced study, Ministry of Human Resource Development (MHRD), and the DBT IISc Partnership Program. The funders had no role in study design, data collection and interpretation, or the decision to submit the work for publication.

## Abbreviations

YSD: Yeast surface display
SSM: Site saturation mutagenesis
FACS: Fluorescence-activated cell sorting
DFE: Distribution of fitness effects
RBD: Receptor binding domain
SARS-CoV-2: Severe acute respiratory syndrome coronavirus 2
ACE-2: Angiotensin-converting enzyme 2

