## Supplementary Figure S1-S13, Supplementary Table S1-S3 for "Prediction of residue-specific contributions to binding and thermal stability using yeast surface display"

**Supplementary materials**

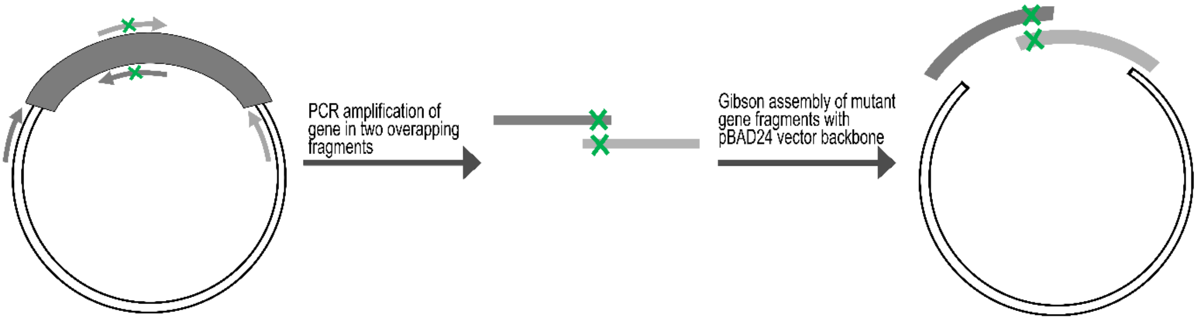


**Supplementary Figure S1: Schematic representation of mutagenesis methodology used to introduce individual mutations in the *E.coli* expressed *ccdB* gene.** The *ccdB* gene was amplified in two fragments using two sets of primers shown in light and dark grey. The location of the mutation is indicated by a green 'X'. The PCR amplified products were gel extracted and 3 fragment Gibson assembly was performed using NdeI and HindIII digested pBAD24 plasmid. The assembled products were transformed in *E.coli* Top10 GyrA cells and individual mutations were confirmed by Sanger sequencing.

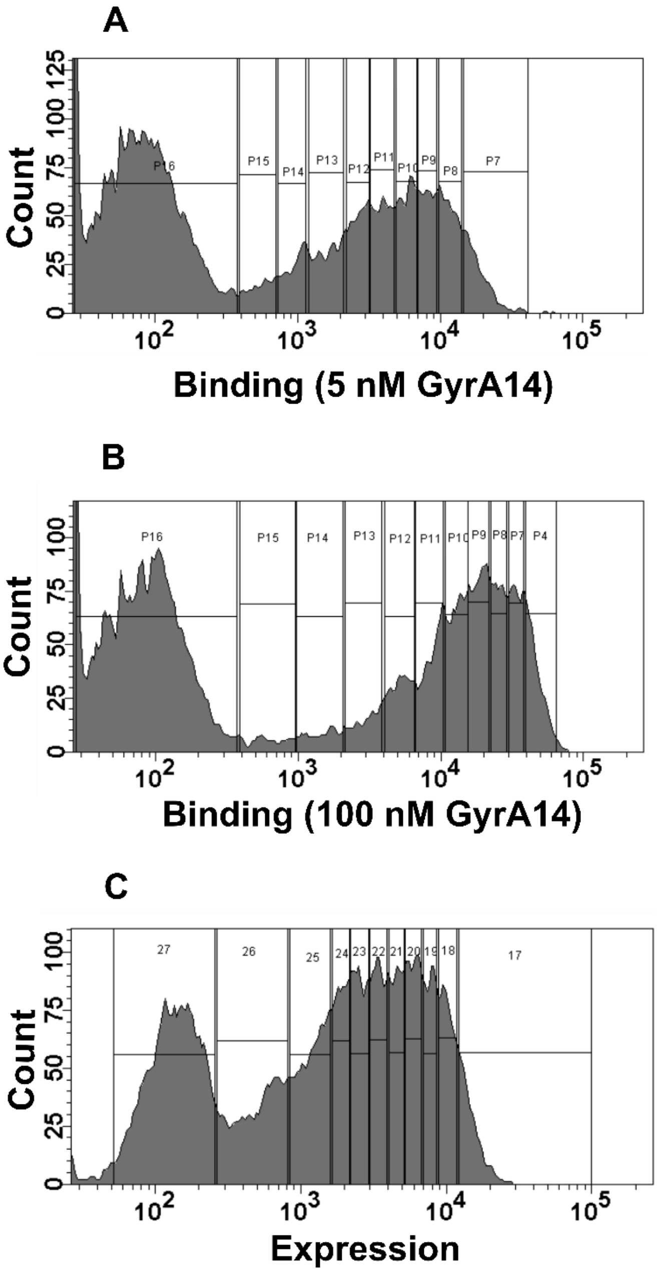


**Supplementary Figure S2: FACS histograms for CcdB mutant library displayed on the yeast cell surface.** Multiple vertical bins were used to sort the library according to (A) binding to 5 nM GyrA14, (B) binding to 100 nM GyrA14 and (C) expression of CcdB mutants. Each bin contained approximately equal numbers of cells.

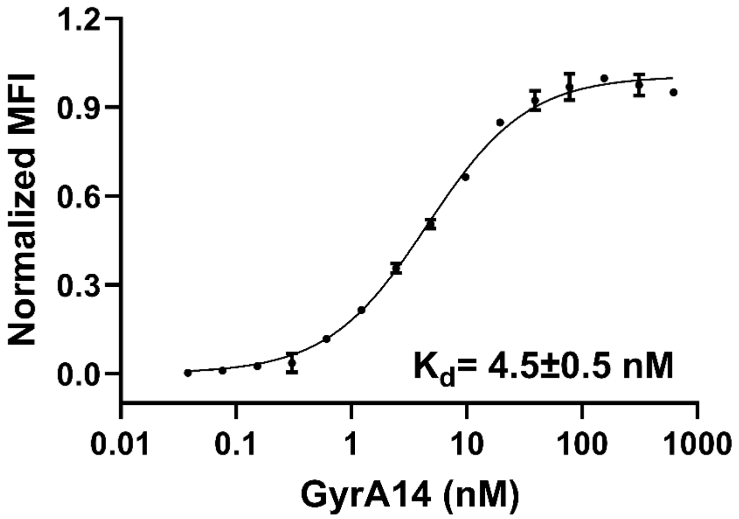


**Supplementary Figure S3: Apparent $K_d$ of interaction between CcdB and GyrA14 on yeast cell surface.** WT CcdB protein was expressed on the yeast cell surface and MFI bind as a function of GyrA14 concentration was used to estimate the $K_d$. Error bars are from two biological replicates.

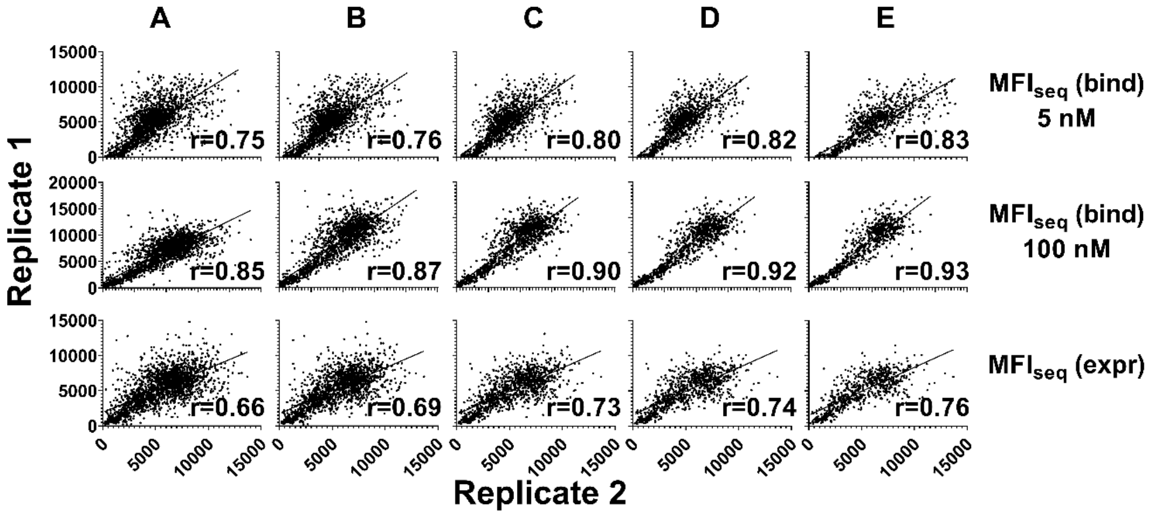


**Supplementary Figure S4: Correlation of $MFI_{seq}$ values between biological replicates obtained at different stringencies (sum of total reads of a mutant across all the bins of binding or expression histogram).** The $MFI_{seq}$ was calculated and a correlation between replicates was measured at stringencies of (A) 25 reads, (B) 50 reads, (C) 100 reads, (D) 150 reads, (E) 200 reads. Only mutants whose total read count exceeded the indicated stringency value in both replicates were considered for analysis. A read count of 50 was chosen for subsequent analyses, as increasing stringency further did not significantly improve the correlation and resulted in fewer mutants being selected.

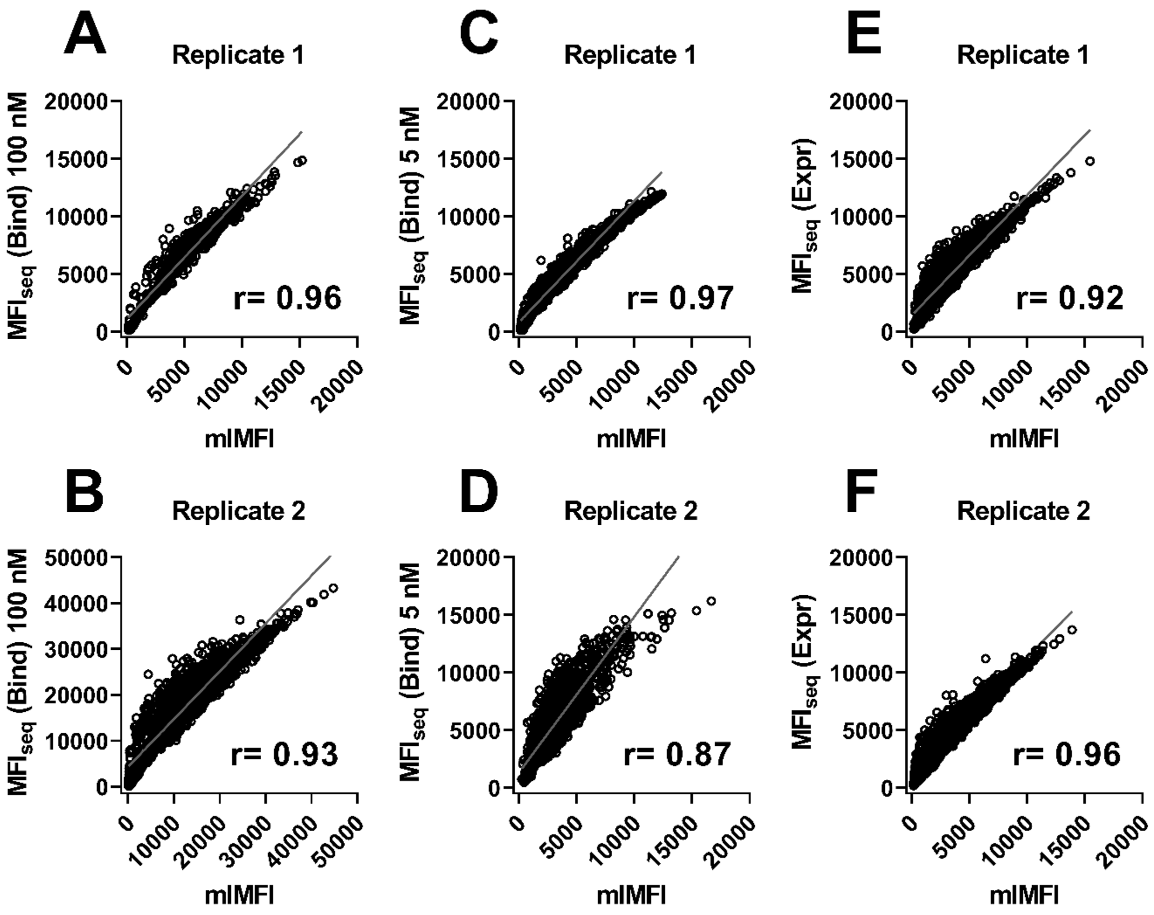


**Supplementary Figure S5: Correlation between MFI of mutants calculated using Maximum Likelihood Method mlMFI using fitdistrplus R package and $MFI_{seq}$.** A good correlation was found between mlMFI and $MFI_{seq}$ for biological replicates at GyrA14 concentrations of 100 nM (A, B), 5 nM (C, D) and for cell surface expression (E, F). Using $MFI_{seq}$ methodology we could calculate the MFI of binding for highly destabilizing as well as nonsense mutants, whereas with mlMFI, for several of these mutants it was not possible to calculate the MFI as all the reads were found in a single bin with the lowest MFI.

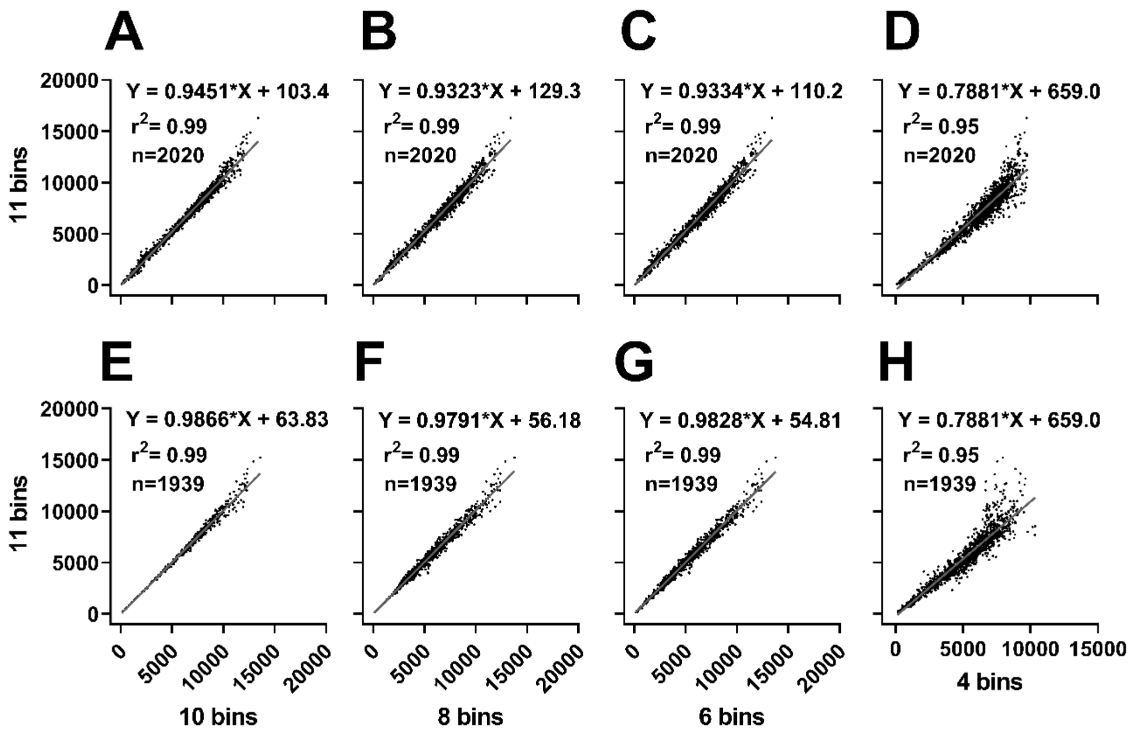


**Supplementary Figure S6: Effect of number of bins on calculated MFI values.** The $MFI_{seq}$ (A-D) and mlMFI (E-H) of mutants was calculated using varying numbers of bins. The binding MFI of CcdB mutants was calculated using 11 bins (y-axis, all panels) and correlated with MFI calculated using 10 bins (A, E), 8 bins (B, F), 6 bins (C, G), 4 bins (D, H). Calculated values for 10, 8 and 6 bins are similar to those with 11 bins for both $MFI_{seq}$ and mlMFI values.

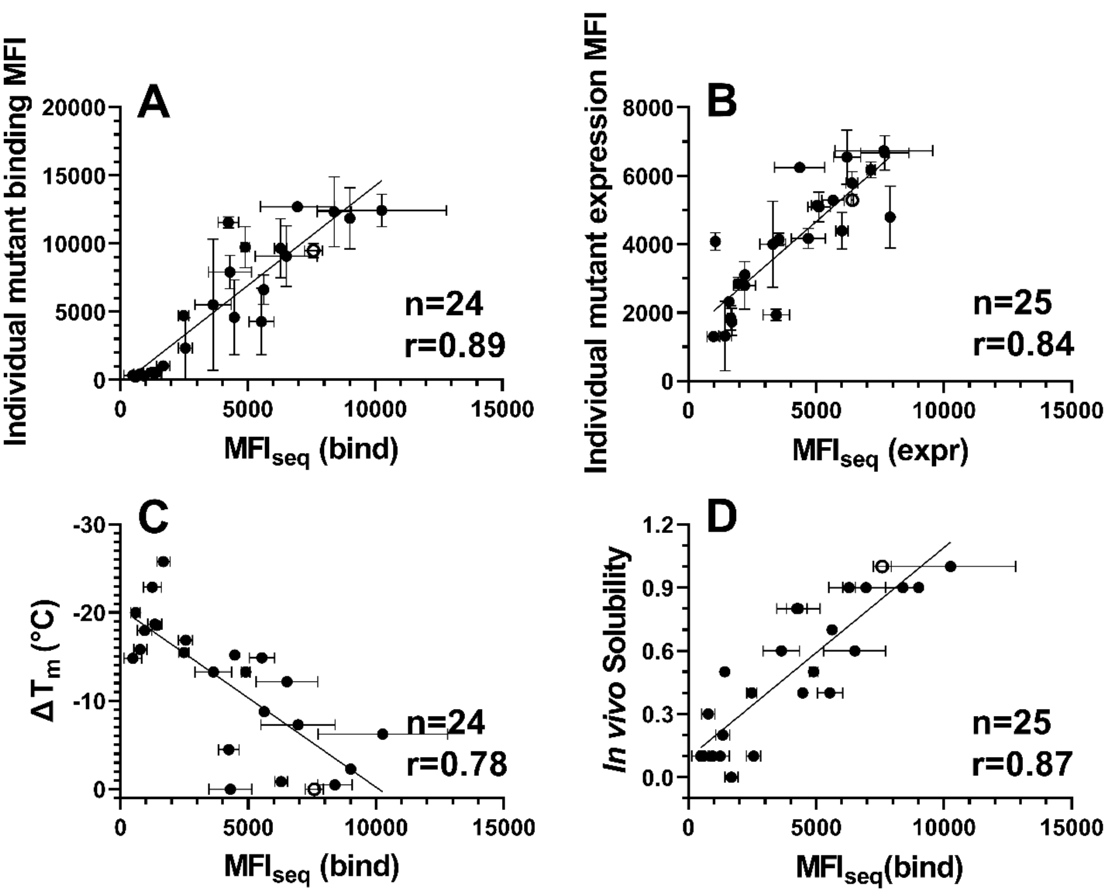


**Supplementary Figure S7: $MFI_{seq}$ correlates well with the MFI of individual mutants.** Correlation between experimentally measured binding MFI from two biological replicates of individually analysed mutants and $MFI_{seq.}$ (A) $MFI_{seq}$ (bind), (B) $MFI_{seq}$ (expr). Correlation of $MFI_{seq}$ (bind) with (C) *in vitro* thermal stability, (D) fractional *in vivo* solubility when expressed in *E.coli.* WT data is shown in open circles.

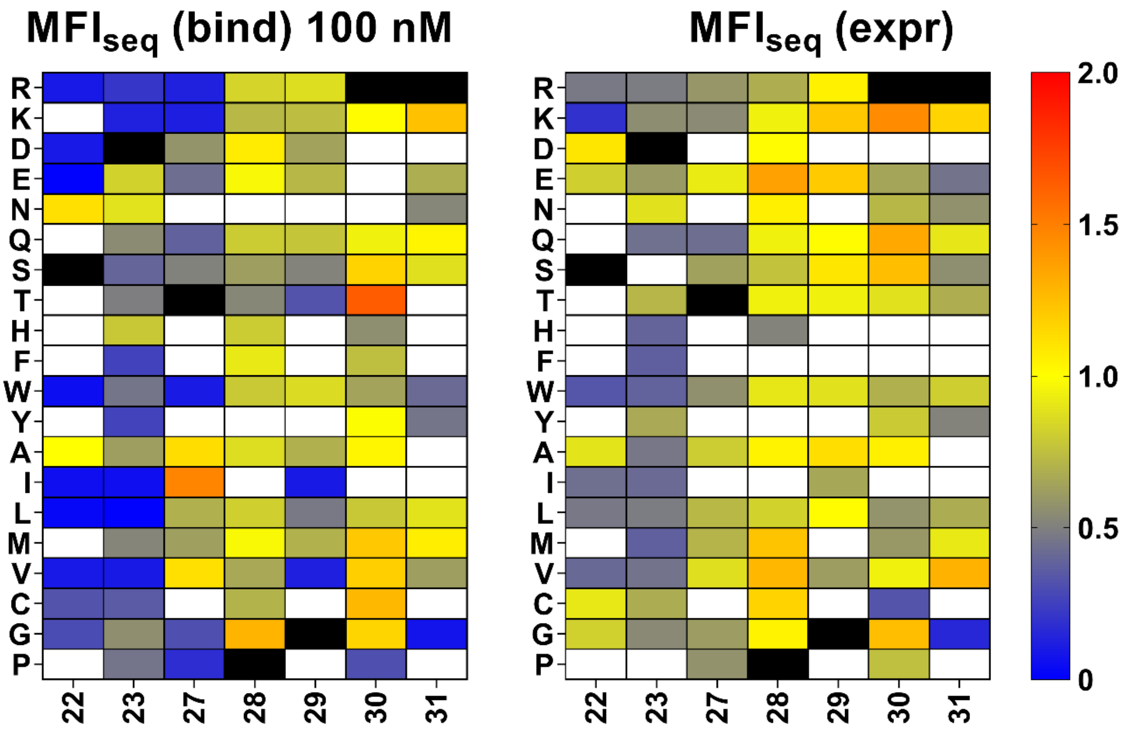


**Supplementary Figure S8: Heat map of normalized binding and expression MFIs of GyrA14 non-interacting residues in the loop connecting β strands S2 and S3.** Blue to red colour represents increasing normalized MFI$_{seq}$ values, black represents the WT residue at the corresponding position. The loop residues 22, 23 and 27 have very high mutational sensitivity in the case of binding at 100 nM of GyrA14. However, expression is largely unaffected by mutation, indicating this region is important for CcdB interaction with GyrA14 even though there is no direct contact with GyrA14 at any of these residues.

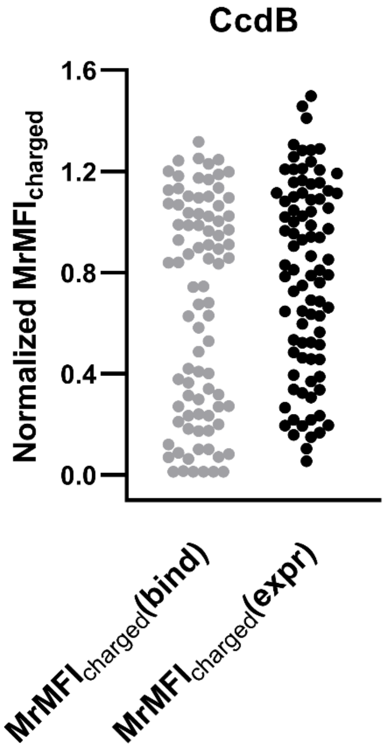


**Supplementary Figure S9: Distribution of MrMFI$_{charged}$ for CcdB.** Binding as well as expression MrMFI$_{charged}$ shows a bimodal distribution in CcdB. k-means clustering was used to calculate the mean and standard deviation of both distributions to differentiate buried and active-site residues.

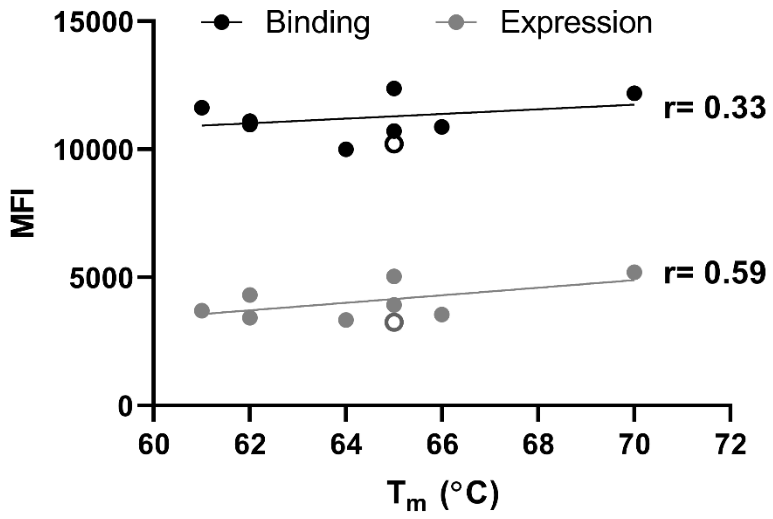


**Supplementary Figure S10: Moderate correlation of MFI values with $T_m$ for mutants with high values of both $MFI_{seq}$ (bind) and $MFI_{seq}$ (expr)**. WT is shown in open circles.

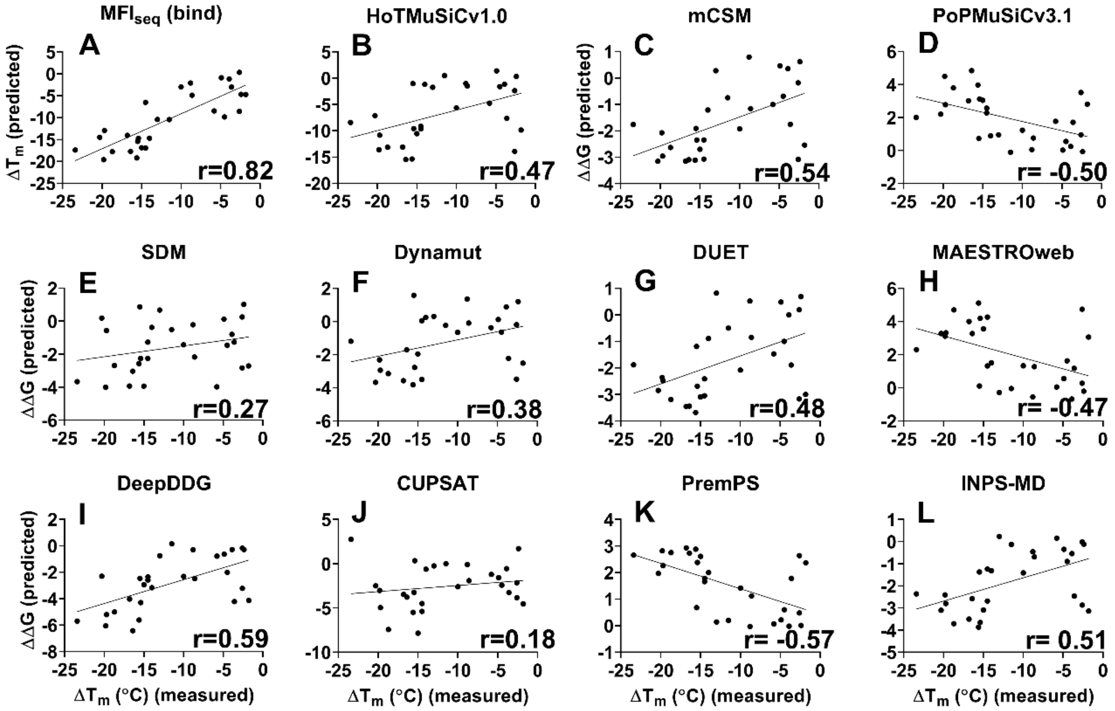


**Supplementary Figure S11: Correlation between predicted and observed $\Delta T_m$ of CcdB mutants.** The stability of CcdB mutants relative to WT was predicted by either (A) $MFI_{seq}$ (bind) or (B-L) existing *in silico* tools. The best correlation was observed between $MFI_{seq}$ (bind) predicted $T_m$ and experimentally measured stability. Thermal stability predictions by *in silico* methods were best for DeepDDG, PremPS and mCSM. For plots with a positive slope, mutants with $\Delta\Delta G>0$ are predicted to be stabilizing, for plots with a negative slope, mutants with $\Delta\Delta G<0$ are predicted to be stabilizing.

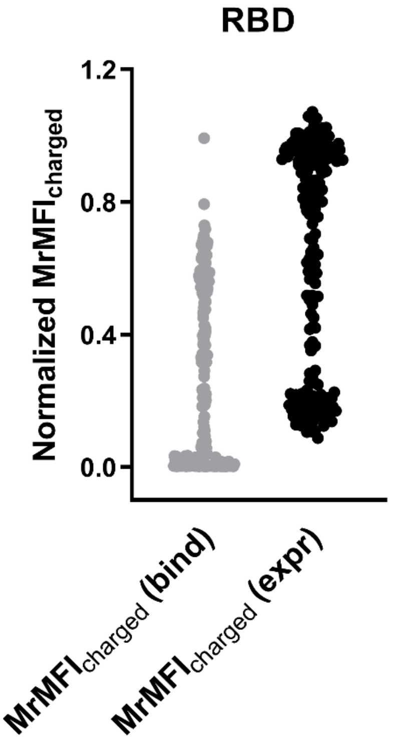


**Supplementary Figure S12: Distribution of MrMFI$_{charged}$ of SARS-CoV-2 RBD.** Normalized MrMFI$_{charged}$ for both expression and binding show a bimodal distribution similar to CcdB. k-means clustering was used to calculate the mean and standard deviation of both distributions to differentiate buried and active-site residues.

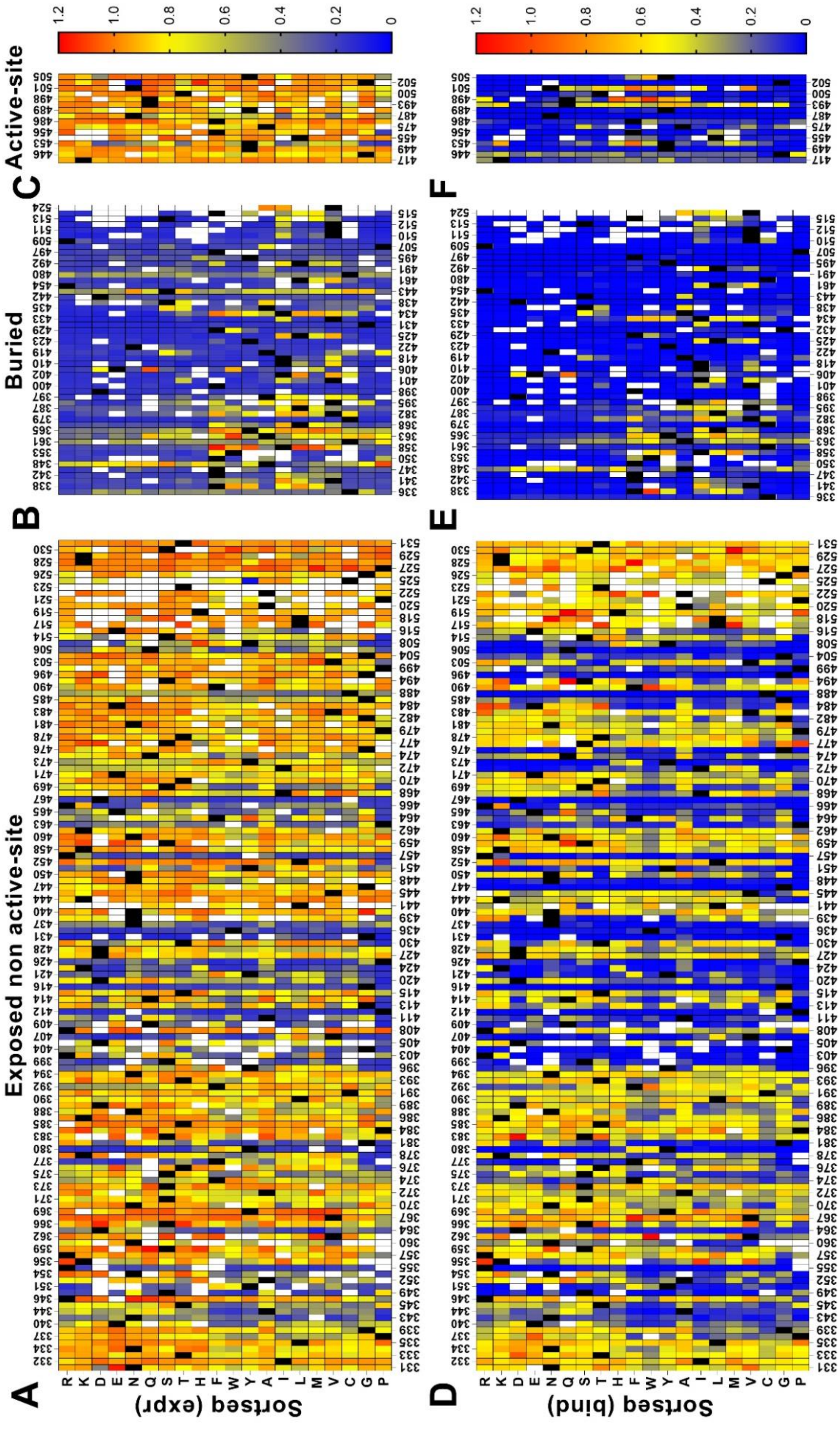


**Supplementary Figure S13: Heatmap of normalized Sortseq MFI values for RBD mutants.** Sortseq (expr) for (A) exposed non active-site residues, (B) buried residues and (C)

active-site residues. Sortseq (bind) at 100 nM ACE-2 for (D) exposed non active-site, (E) buried and (F) active-site residues. Exposed, buried (PDB ID:7KMH) and active-site (PDB ID:6M0J) residues are segregated based on the crystal structure. Blue to red colour represents increasing normalized Sortseq MFI values, black colour shows the WT residue at the corresponding position. White colour indicates that the mutant is not available.

**Tables**

**Supplementary Table S1:** Correlation coefficient (r) for $MFI_{seq}$ values of individual mutants between two replicates at different stringencies. For stringencies of 50 reads or greater there is good correlation of both $MFI_{seq}$ (bind) and $MFI_{seq}$ (expr) between replicates.

| Stringency | Experiment 1 | Experiment 2 | r |
|---|---|---|---|
| 25 reads | $MFI_{seq}$ (expr) | $MFI_{seq}$ (expr) | 0.66 |
| | $MFI_{seq}$ (bind) at 5 nM GyrA14 | $MFI_{seq}$ (bind) at 5 nM GyrA14 | 0.75 |
| | $MFI_{seq}$ (bind) at 100 nM GyrA14 | $MFI_{seq}$ (bind) at 100 nM GyrA14 | 0.85 |
| 50 reads | $MFI_{seq}$ (expr) | $MFI_{seq}$ (expr) | 0.69 |
| | $MFI_{seq}$ (bind) at 5 nM GyrA14 | $MFI_{seq}$ (bind) at 5 nM GyrA14 | 0.76 |
| | $MFI_{seq}$ (bind) at 100 nM GyrA14 | $MFI_{seq}$ (bind) at 100 nM GyrA14 | 0.87 |
| 100 reads | $MFI_{seq}$ (expr) | $MFI_{seq}$ (expr) | 0.73 |
| | $MFI_{seq}$ (bind) at 5 nM GyrA14 | $MFI_{seq}$ (bind) at 5 nM GyrA14 | 0.80 |
| | $MFI_{seq}$ (bind) at 100 nM GyrA14 | $MFI_{seq}$ (bind) at 100 nM GyrA14 | 0.90 |
| 150 reads | $MFI_{seq}$ (expr) | $MFI_{seq}$ (expr) | 0.74 |
| | $MFI_{seq}$ (bind) at 5 nM GyrA14 | $MFI_{seq}$ (bind) at 5 nM GyrA14 | 0.82 |
| | $MFI_{seq}$ (bind) at 100 nM GyrA14 | $MFI_{seq}$ (bind) at 100 nM GyrA14 | 0.92 |
| 200 reads | $MFI_{seq}$ (expr) | $MFI_{seq}$ (expr) | 0.76 |
| | $MFI_{seq}$ (bind) at 5 nM GyrA14 | $MFI_{seq}$ (bind) at 5 nM GyrA14 | 0.83 |
| | $MFI_{seq}$ (bind) at 100 nM GyrA14 | $MFI_{seq}$ (bind) at 100 nM GyrA14 | 0.93 |

**Supplementary Table S2:** Correlation coefficient (r) for $MFI_{seq}$ values of individual mutants with CcdB structural parameters calculated from the WT structure (PDB ID: 3VUB). All correlations are improved when $MFI_{seq}$ values are averaged (MrMFI) over all mutants at the given residue.

| Parameter 1 | Parameter 2 | r |
|---|---|---|
| Depth | $MFI_{seq}$ (bind)at 5 nM GyrA14 | -0.29 |
| | MrMFI (bind)at 5 nM GyrA14 | -0.33 |
| Depth | $MFI_{seq}$ (bind) at 100 nM GyrA14 | -0.32 |
| | MrMFI (bind) at 100 nM GyrA14 | -0.44 |
| Depth | $MFI_{seq}$ (expr) | -0.45 |
| | MrMFI (expr) | -0.58 |
| % Accessibility | $MFI_{seq}$ (bind)at 5 nM GyrA14 | 0.27 |
| | MrMFI (bind) at 5 nM GyrA14 | 0.30 |
| % Accessibility | $MFI_{seq}$ (bind) at 100 nM GyrA14 | 0.27 |
| | MrMFI (bind) at 100 nM GyrA14 | 0.33 |
| % Accessibility | $MFI_{seq}$ (expr) | 0.37 |
| | MrMFI (expr) | 0.44 |
| RankScore | $MFI_{seq}$ (bind)at 5 nM GyrA14 | -0.37 |
| | MrMFI (bind) at 5 nM GyrA14 | -0.37 |
| RankScore | $MFI_{seq}$ (bind)at 100 nM GyrA14 | -0.47 |
| | MrMFI (bind) at 100 nM GyrA14 | -0.61 |
| RankScore | $MFI_{seq}$ (expr) | -0.53 |
| | MrMFI (expr) | -0.57 |

**Supplementary Table S3**

The specificity, sensitivity and accuracy of exposed non active-site, buried site and exposed active-site residue identification solely from $MFI_{seq}$ or Sortseq.

| | **Specificity**[a] | | **Sensitivity**[a] | | **Accuracy**[a] | |
|---|---|---|---|---|---|---|
| **Proteins** | CcdB | RBD | CcdB | RBD | CcdB | RBD |
| **Exposed non-active site** | 0.98 | 0.84 | 0.82 | 0.74 | 0.92 | 0.80 |
| **Buried** | 1 | 0.87 | 0.76[b] | 0.70[b] | 0.93 | 0.81 |
| **Exposed active-site** | 0.97 | 0.98 | 0.88 | 0.78 | 0.95 | 0.96 |

[a]The specificity, sensitivity and accuracy were calculated using the following formulae, where TN, TP, FN and FP correspond to true negative, true positive, false negative and false positive respectively.

Specificity = $\frac{TN}{TN+FP}$

Sensitivity = $\frac{TP}{TP+FN}$

Accuracy = $\frac{TP+TN}{TP+TN+FP+FN}$

[b]Sensitivity increases to 0.79 and 0.87 for CcdB and RBD respectively if false negative Gly residues are not considered.
